# The methyl-CpG-binding protein 2 inhibits cGAS-associated signaling

**DOI:** 10.1101/2025.01.30.635818

**Authors:** Hanane Chamma, Soumyabrata Guha, Moritz Schüssler, Yasmine Messaoud-Nacer, Pierre Le Hars, Mohammad Salma, Joe McKellar, Joanna Re, Morgane Chemarin, Arnaud Carrier, Michael A. Disyak, Clara Taffoni, Robin Charpentier, Zoé Husson, Emmanuel Valjent, Charlotte Andrieu-Soler, Eric Soler, Karim Majzoub, Isabelle K. Vila, Nadine Laguette

## Abstract

The detection of cytosolic dsDNA is tightly regulated to avoid pathological inflammatory responses. A major pathway involved in their detection relies on the cyclic GMP-AMP synthase (cGAS) that triggers activation of the Stimulator of interferon genes (STING) which subsequently drives the expression of inflammatory genes and type I Interferons (IFNs). Here, we show that the methyl-CpG-binding protein 2 (MECP2), a major transcriptional regulator, controls dsDNA-associated inflammatory responses. We show that the presence of cytosolic dsDNA promotes MECP2 export from the nucleus to the cytosol where it interacts with dsDNA, dampening cGAS activation. Our data also indicate that MECP2 export from the nucleus partially phenocopies MECP2 deficiency, leading to the expression of inflammatory and interferon stimulated genes, enforcing an antiviral state. Finally, we also show that MECP2 displacement from the nucleus following dsDNA stimulation is sufficient to disrupt its canonical function, leading to the reactivation of otherwise repressed genes, such endogenous retroelements of the Long interspersed nuclear element-1 (LINE-1) family. Re-expression of the latter led to the accumulation of DNA species feeding cGAS-dependent signaling and can be dampened by reverse transcriptase inhibitors. We thus establish a previously unforeseen direct role of MECP2 in the regulation of the breadth and nature of dsDNA-associated inflammatory responses. Furthermore, our results suggest that targeting dsDNA-associated pathways or pharmacological inhibition of LINE-1 may bear therapeutic hopes for Rett syndrome (RTT) patients that present with MECP2 deficiency.

## Introduction

Cytosolic dsDNAs are recognized drivers of chronic inflammation in a broad range of human diseases ^1^. A major pathway involved in detecting cytosolic dsDNA and triggering the subsequent activation of inflammatory responses, relies on the cyclic GMP-AMP (cGAMP) synthase (cGAS) pathogen recognition receptor ^2–5^. The interaction of cytosolic dsDNAs with cGAS leads to the production of the 2’3’-cGAMP second messenger, which in turn interacts with the stimulator of interferon genes (STING) adaptor protein, promoting the recruitment and activation of transcription factors, such as the Interferon regulatory factor 3 (IRF3), that drive a transcriptional program ultimately leading to the production of type I Interferons (IFN) and inflammatory cytokines^2–5^. Regulation of the cGAS-STING signalling axis at several levels has been reported, including STING degradation ^6^, or regulators of cGAS catalytic activity ^7,8^. In contrast, although the detection of dsDNA by cGAS being a key rate-limiting step in the activation of cGAS-STING signalling, few regulators of this interaction have been identified to date ^9^. Those include the Three-prime exonuclease 1 (Trex1), the ablation of which feeding cGAS-STING-dependent signaling through cytosolic DNA accumulation, and underlying the most severe form of Aicardi-Gouttière Syndrome (AGS) ^10^. This highlights the importance of the regulation of the cytosolic DNA-cGAS interaction for the regulation of pro-inflammatory signalling.

Pathological activation of cGAS-STING signalling has been documented in a wide-range of neurological disorders ^11^, mostly highlighting a link between chronic STING signalling and neurodegeneration ^12–14^. Similarly, chronic inflammatory responses and immunological dysfunction have been reported in the Rett syndrome neurodevelopmental disorder (RTT; OMIM 312750). RTT is a rare genetic disorder frequently associated with *de novo* mutations in the *Methyl-CpG-binding protein 2* (*MECP2*) X-linked gene, giving rise to dysfunctional MECP2 ^15^. Neurological manifestations of RTT have been shown to be reversible ^16^ and accompanied by immunological dysfunction as well as chronic low-grade inflammation ^17–19^. Indeed, absence of MECP2 sensitizes immune cells to pro-inflammatory stimulation ^20^. While inflammation has been proposed to be associated with RTT disease progression and severity ^21,22^, there is no mechanism explaining the onset of chronic inflammation in RTT.

MECP2 is a broadly expressed nuclear protein known to operate as a major transcription regulator ^23^. MECP2 was initially described to bind and repress the expression of methylated regions of the genome ^23,24^, but the functions of MECP2 in the regulation of the mammalian genome have expanded in recent years. Indeed, MECP2 is now recognized to play context-dependent roles in gene expression and chromatin architecture regulation through DNA methylation-independent mechanisms ^25–27^. MECP2 was shown to associate with and repress genomic locations corresponding to endogenous retroelements, such as the Long interspersed nuclear element-1 (LINE-1) ^28^. Interestingly, increased LINE-1 activity has been shown to be sufficient to generate DNA substrates for cGAS-STING activation ^29^.

Altogether, the current state-of-the-art suggests a link between MECP2 and dsDNA-mediated activation of the cGAS-STING pathway. Thus, we here investigated the potential role of MECP2 as a regulator of dsDNA-associated inflammatory responses.

## Materials and Methods

### Mecp2-deficient mouse models

*Post mortem* samples from Mecp2-deficient (Jackson (B6.129P2(c)- Mecp^2tm1–1Bird^) male mice were a kind gift from Emmanuel Valjent (IGF Montpellier) and Adrian Bird (University of Edinburgh). Meta-analyzed mouse mRNA sequencing data ^30^ where from hippocampal mRNA profiles of 3-month-old mice following delivery of either a control shRNA sequence or a MeCP2-specific shRNA.

### Cells and Cell Culture

Wild-type (WT) murine embryonic fibroblast (MEF) and cGAS-deficient MEF (MEF*^cGas-/-^*) were a gift of S. R. Paludan. BHK21 cells were a gift of Olivier Moncorgé (IRIM, Montpellier). Parental RAW246.7 (*Mus musculus*), NIH3T3 (*Mus musculus*), HEK293T (*Homo sapiens*) and THP-1 (*Homo sapiens*) cells were obtained from the American Type Culture Collection (ATCC). MEF^gMecp2^ and RAW246.7 ^gMecp2^ cell lines, along with their respective controls, were generated in the laboratory using the CRISPR-Cas9 technology. MEF, HEK293T and RAW246.7 cells were maintained in Dulbecco’s Modified Eagle Medium (DMEM) supplemented with 10 % Fetal Bovine Serum (FBS, Eurobio), 1% L-glutamine (Lonza), 1% Penicillin/Streptomycin (Lonza). In-house generated knockout cell lines were maintained in the presence of the puromycin selection antibiotic. THP-1 cells were cultured in Roswell Park Memorial Institute (RPMI, Lonza) supplemented with 10% FBS, 1% penicillin/streptomycin and 1% L-glutamine. All cell lines were maintained at 37 °C, under 5% CO_2_.

### Plasmids, constructs and synthetic nucleic acid probes

MEF ^gMecp2^ and RAW264.7 ^gMecp2^ and control cell lines, were generated by lentiviral transduction followed by selection, using the LentiCRISPRv2puro plasmid (Addgene #82416) in which control non-targeting or Mecp2-targeting guide RNAs (gRNA) were cloned using the following sequences:

#### Control non-targeting gRNA

Forward (F): CACCGACGGAGGCTAAGCGTCGCAA;
Reverse (R): AAACTTGCGACGCTTAGCCTCCGTC

#### Mecp2-targeting gRNA

F: CACCGCGCTCCATTATCCGTGACCG;
R: CGCGAGGTAATAGGCACTGGCCAAA

To generate the Flag-tagged Mecp2 expressing construct, the Mecp2 gene was amplified by PCR from peGFP-N1-Mecp2 WT plasmid (Addgene #110186) using the Phusion High-Fidelity DNA polymerase kit (M0530L) followed by cloning into the pOZ vector ^31^.

Synthetic nucleic acid probes used for *in vitro* and in-cell pulldowns were purchased as ssDNA 80 base pair (bp)-long ssDNA probes, bearing or not 5’ biotin or Cyanin 3-labels, from Integrated DNA Technologies (IDT). The 5’ biotin and Cyanin 3 tags were always on the sense strand. When dsDNA probes were used, they were generated by annealing of sense and anti-sense ssDNA probes and integrity assessed as described in ^32^. Sequences of nucleic acid probes used in the present study are:

#### Sense (S) ssDNA (S-ssDNA)

ACATCTAGTACATGTCTAGTCAGTATCTAGTGATTATCTAGACATACATGATCTATGACATATATAGTGGATAAGTGTGG.

#### Anti-sense (AS) ssDNA (AS-ssDNA)

CCACACTTATCCACTATATATGTCATAGATCATGTATGTCTAGATAATCACTAGATACTGACTAGACATGTACTAGATGT.

More precisely, dsDNA was obtained by annealing of S-ssDNA and AS-ssDNA; 5’biotin-bearing dsDNA (b-dsDNA) were obtained by annealing 5’-biotinylated S-ssDNA and AS-ssDNA;; and Cy3-labelled dsDNA (Cy3-dsDNA) were obtained by annealing Cyanin3-labelled S-ssDNA and AS-ssDNA.

### Generation of knock-out cell lines

Lentiviral particles used to generate MEF^gMecp2^, RAW264.7^gMecp2^ and control cell lines were produced by co-transfection of 2 × 10^6^ HEK293T with 5 μg of LentiCRISPRv2puro plasmid expressing non-targeting or Mecp2-targeting gRNAs, together with 5 μg of psPAX2, and 1 μg of pMD2.G, using the calcium phosphate transfection protocol. Lentiviral particles were harvested 48h after transfection and filtered (0.45 µM) prior to transduction of MEF or RAW264.7. Selection was initiated 72 h post transduction using 1.5 μg/ml or 4 of μg/ml puromycin for MEFs or RAW264.7, respectively. Protein levels of Mecp2 were controlled by Western blot (WB) after 3 days of selection.

### Transfection and treatments

For gene expression analyses, cells were transfected or not with 0.2 or 2 μg of ssDNA or dsDNA per well of 6-well plates. For in-cell pulldowns, cells were transfected or not with 1 or 10 µg of b-ssDNA, or b-dsDNA when experiment were conducted in 10 cm dishes. Two or 20 µg of nucleic acid probes were transfected when experiments were conducted in 15 cm dishes. Transfections were performed using JetPrime (Polyplus) in DMEM or Opti-MEM (GIBCO) for 1, 3, and/or 6 hours as indicated in the figure legends.

For nuclear export inhibition, cells were treated with 20 nM of Leptomycin B (LMB, Cell signaling technology (CST)) for 1 hour prior to dsDNA transfection for 3 hours. For STING inhibition, cells were treated with 1 μM of H-151 for 1 hour prior to dsDNA transfections for 6 hours. Reverse transcriptase activity was inhibited by treating cells with 25 μM of Tenofovir for 24 hours prior to dsDNA transfection for 6 or 24 hours.

Following transfection or treatment, cells were collected for gene expression analysis, WB, in-cell pulldowns, immunoprecipitations or cGAMP quantification, or fixed for immunofluorescence.

### RNA extraction and gene expression analyses

Total RNA was isolated with TRIzol reagent (Invitrogen) or using the GenElute^TM^ Mammalian total RNA kit (Sigma #SLBW4972). RNA concentration was measured with a Nanodrop spectrophotometer (ND-1000, Nanodrop Technologies), prior to treatement with TURBO DNase (Ambion) and cDNA synthesis from 1-2 μg RNA using SuperScript IV (Invitrogen) using Oligo(dT) and quantification with a LightCycler 480 cycler (Roche) using SYBR Green Master Mix (Takara) and appropriate primers. Relative quantities of the transcript were calculated using the ΔΔCt method, using the heat shock protein 90 (*Hsp90*) or Glyceraldehyde3-phosphate dehydrogenase (*Gapdh*) for normalization. The RT-qPCR primers used are listed below:

*<u>Ccl5:</u>* **F**: <u>CAGCAAGTGCTCCAATCTTGC</u>; **R**: <u>CCACTTCTTCTCTGGGTTGGC</u>
*<u>Cxcl10</u>*: **F:** ATGACGGGCCAGTGAGAATG; **R**: TCAACACGTGGGCAGGATAG
*<u>Gapdh</u>*: **F**: TTCACCACCATGGAGAAGGC; **R**: GGCATCGACTGTGGTCATGA
*<u>Hsp90</u>*: **F**: GTCCGCCGTGTGTTCATCAT; **R**: GCACTTCTTGACGATGTTCTTGC
*<u>Ifnβ</u>*: **F:** CTGCGTTCCTGCTGTGCTTCTCCA; **R**: TTCTCCGTCATCTCCATAGGGATC
*<u>Il6</u>*: **F**: GACTTCCATCCAGTTGCCTTCT; **R**: TCCTCTCCGGACTTGTGAAGTA
*<u>Isg15</u>*: **F**: GTGCTCCAGGACGGTCTTAC; **R**: CTCGCTGCAGTTCTGTACCA
*<u>Mecp2</u>*: **F**: TCAGAAAGCTCAGGCTCTGC; **R**: CCCGGTCACGGATAATGGAG
*<u>Mx2:</u>* **F**: CCTATTCACCAGGCTCCGAA; **R**: CGTCCACGGTACTGCTTTTC
*<u>Oas1</u>*: **F**: TGCATCAGGAGGTGGAGTTTG; **R**: ATAGATTCTGGGATCAGGCTTGC
*<u>Oas1b</u>*: **F**: GCAAAGGCACCACACTCAAG; **R**: CTCTCATGCTGAACCTCGCA
*<u>Oasl1:</u>* **F**: CAGGAGCACTACAGACGTGG; **R**: GGTTACTGAGCCCAAGGTCC
*<u>Oas2:</u>* **F**: GAGTGGGAGGTGACGTTTGA; **R**: GAGTGGGAGGTGACGTTTGA
*<u>Oas3:</u>* **F**: CCAAAGCGTGGACTTTGACG; **R**: GCAGCTCTGTGAAGCAGGTA
*<u>Slfn5:</u>* **F**: CGAAATCATCTCGCAAGCCG; **R**: TGGTGGCAGATTCAAGCCAA

### Whole cell lysate preparation

For WB, pulldowns and immunoprecipitations, cells were harvested on ice in cold phosphate-buffered saline (PBS) using a cell scraper and lysed in 5 packed cell volume (PCV) of TENTG-150 [20 mM tris-HCl (pH 7.4), 0.5 mM EDTA, 150 mM NaCl, 10 mM KCl, 0.5% Triton X-100, 1.5 mM MgCl_2_, and 10% glycerol], supplemented with 10 mM β-Mercaptoethanol, 0.5 mM phenylmethylsulfonyl fluoride (PMSF) and phosphatase inhibitor (PhosphoSTOP, Sigma-Aldrich) for 30 min at 4°C. Cell lysates were centrifuged at 12,000g for 30 min at 4°C and supernatants collected. Protein concentration was determined using the Bradford assay (Sigma-Aldrich).

### Nuclear-cytoplasmic fractionation

For subcellular fractionation, cells were harvested and pellet size normalized prior to lysis in 5 PCV of low Salt buffer [100 mM NaCl, 0.1% Triton, 20 mM Tris pH 7.4, 2 mM MgCl2, 0.5 mM EDTA, and 10% Glycerol] extemporaneously supplemented with 0.2 mM PMSF and 10 mM β-Mercaptoethanol for 20 min on wheel at 4°C. Nuclei were pelleted at 2,000 g for 10 min and cytosolic extracts were collected in a fresh tube. Nuclei were then lysed by adding 5 PCV high Salt buffer [340 mM NaCl, 0.1% Triton, 20 mM Tris pH 7.4, 2 mM MgCl_2_, 0.5 mM EDTA, and 10% Glycerol] extemporaneously supplemented with 0.2 mM PMSF and 10 mM β-Mercaptoethanol for 30 min on a wheel at 4°C. Soluble nuclear extracts were collected after centrifuging the samples at 12,000g for 10 min. When the Dignam nuclear and S100 extracts were used, they were prepared according to the protocol described in ^31^.

### Western Blot analyses

Protein samples were prepared in Laemmli buffer and heated at 95°C for 5 min prior to resolution by sodium dodecyl sulfate–polyacrylamide gel electrophoresis (SDS-PAGE) using precast 10% or 12% gels (Invitrogen Novex Tris-glycine) followed by transfer onto nitrocellulose membranes using the Trans-Blot Turbo Transfer System (Biorad). Proteins were visualized on membranes using Ponceau S solution (Sigma-Aldrich) prior to 30 min blocking with (PBS) containing 0.1% Tween (PBS-T) supplemented with 5% milk, and incubation overnight at 4°C for phosphorylated targets or 90 min at room temperature with primary antibodies in 5% milk/PBS-T or 5%BSA/PBS-T. Primary antibodies used include: anti-Mecp2 (1:500; Cell signaling D4F3), mouse-specific anti-cGas (1:1000; Cell signaling D3080), anti-phosphorylated Irf3 (1:500; Cell Signaling 4D4G), anti-Irf3 (1:1000; Cell Signaling D83B9), anti-phosphorylated Tbk1 (1:1000; Cell Signaling D52C2), anti-Tbk1 (1:1000; Cell Signaling D1B4), anti-Sting (1:1000; Cell Signaling D2P2F), anti-phosphorylated Sting (1:1000; Cell Signaling D8F4W), anti-Hsp90 (1:1000; Cell Signaling C45G5), anti-Gapdh (1:5000; Proteintech Europe 60004-1-Ig), anti-Lamin B1 (1:1000, Santa Cruz Biotechnology sc-374015), anti-Acetylated Histone H3 (1:1000; Santa Cruz Biotechnology sc-56616), anti-Flag (1:1000; Sigma F1804), anti-Tubulin α (66031-1-Ig, Proteintech Europe, 1:10,000), and anti-Ranbp1 (1:100; Santa Cruz Biotechnology #sc-374352). Membranes were incubated with Horseradish peroxidase (HRP)-coupled secondary antibodies (Cell Signaling) at 1:2000 dilution for 1 hour at room temperature. Immunoreactivity was detected by Chemiluminescence (SuperSignal West Pico or Femto Thermo Scientific). Images were acquired on a ChemiDoc (Bio-Rad) or Amersham bioluminescence detection imager.

### *In vitro* biotinylated nucleic acid pull-down

*In vitro* pull-downs were performed using either whole cell lysates or fractionated extracts using 30 μl (10mg/ml) of Dynabeads M280 slurry per condition. Dynabeads were blocked overnight at 4°C under agitation, in blocking buffer [20 mM Hepes pH 7.9, 0.05% NP40, 150 mM NaCl, 15% Glycerol, 2 mM DTT, and 20 mg/ml BSA]. Coupling of beads with 3 μg of biotinylated nucleic acids (b-ssDNA, b-dsDNA or b-4mCGdsDNA) was subsequently performed in annealing buffer [60mM NaCl,10 mM Tris pH 7.5, 0.2mM EDTA) during 15 min at 25°C under gentle agitation. Nucleic acid-coupled beads were then washed twice using washing buffer [1 mM NaCl, 1 mM EDTA, and 5 mM Tris pH 7.4] and equilibrated in TENTG-150. Subsequently, 4 mg of total protein were added to the equilibrated beads and incubated at 4°C for 3 hours in low-binding tubes (Axygen) on wheel. Following incubation, three consecutive washes were performed with TENTG-150 changing tubes at the first and last washes. Bound proteins were eluted in 30 μl of Laemmli buffer for 5min at 95°C prior to WB analyses. The interaction between proteins and tested biotinylated nucleic acids was assessed by western Blot.

### In-cell biotinylated nucleic acid pull-down

Cells were transfected with 1 to 4 μg/ml of biotinylated synthetic nucleic acid probes using JetPrime, for up to 6 hours prior to lysis in TENTG-150 for 30 min at 4°C. Lysates were centrifuged for 30 min at 12,000g at 4°C prior to 45 minutes incubation at 4°C with 30 μl of Dynabeads M280 blocked and equilibrated as above. Following the incubation, three washes were performed as above prior to elution of bound proteins in 30 μl Laemmli buffer and subsequent WB analyses.

### Immunoprecipitation

Endogenous immunoprecipitation was performed from 1mg of total protein from whole cell lysates using 1µg of Mecp2-targeting antibody or control IgG. After an overnight incubation at 4°C on a wheel, immunoprecipitation was performed using Protein G Sepharose beads. After 3 washes in TENTG-150, bound material was eluted in Laemmli buffer prior to WB analyses.

### Immunofluorescence and microscopy analysis

For analysis of Mecp2 subcellular localization following nucleic acid challenge. 3×10^5^ MEFs or 1×10^6^ RAW246.7 cells were seeded onto glass coverslips prior to transfection using either with unlabeled ssDNA or dsDNA, or with 0.5 µg of Cyanin3-labeled dsDNA (Cy3-dsDNA) as described above. Six hours post transfection, cells were fixed at room temperature for 15 min in PBS containing 4% para-formaldehyde (PFA). Following fixation, cells were permeabilized in PBS containing 0.1% Triton X-100 at room temperature prior to blocking in PBS-T supplemented with 5% BSA for 30 min at room temperature. The procedure used for quantification of Mecp2 subcellular localization following infection is described in the corresponding sections.

Coverslips were incubated at 37°C for 45 min with primary antibodies in PBS-T. Primary antibodies used are: anti-Mecp2 (Cell signaling #3456T) used at 1:50 dilution, anti-cGas (Cell signaling #31659) used at 1:100 dilution, anti-IRF3 (Cell signaling #4302S) used at 1:100 dilution, anti-Ranbp1 (Santa Cruz Biotechnology #sc-374352) used at 1:50 dilution, anti-dsDNA (Abcam #ab27156) used at 1:100 dilution, anti-5-hydroxymethyl-cytosine (Cell signaling #51660S) used at 1:400 dilution, and anti-GFP (Abcam #ab290) used at 1:100 dilution. Following 3 washes in PBS-T, coverslips were incubated for 30 min at 37°C with appropriate secondary antibodies at 1:200 dilution in PBS-T. Secondary antibodies used include: Alexa Fluor 488 coupled goat anti-Rabbit IgG, (Thermofischer #R37116), Alexa Fluor 488 goat anti-Mouse IgG, (#A11001, Thermofischer), Alexa Fluor 594-coupled goat anti-Mouse IgG (Thermofischer #R37121), and Alexa Fluor 594-coupled goat anti-Rabbit IgG (#R37117, Thermofischer). After incubation with secondary antibody, coverslips were washed 5 times in PBS-T. Nuclei were stained with 1 µg/ml DAPI in PBS for 5 min. The coverslips were washed 5 times with PBS prior to mounting in anti-fade media (Vectashield). Images were acquired by Apotome microscope (Zeiss) using the ZEN (blue edition) software, with a 63X oil objective, and images were processed with Omero or Fiji software tools.

### cGAMP quantification

Cells were transfected in OptiMEM with dsDNA (2 µg) for 6 hours, harvested in ice-cold PBS, counted for normalization and extracted using the Mammalian Protein Extraction Reagent (M-PER) buffer (ThermoFisher) prior to quantification using the 2’3’-cGAMP enzyme-linked immunosorbent assay (ELISA) (Cayman), according to the manufacturer’s protocol.

### HSV-1 Infection

The HSV-1 KOS-64-GFP strain (HSV-1-GFP) was a gift from S R. Paludan (Aarhus university, Danemark). HSV-1 was amplified on Vero cells, aliquoted and frozen at −80°C. Titration was performed on Vero cells, by serial dilutions and plaque formation assessment to determine the multiplicity of infection (MOI). For gene expression analysis, 2.5×10^5^ cells were seeded per well of 6-well plates 24 hours prior to infection with 5 MOI for 6 or 16 hours. Cells were subsequently harvested and RNA extracted with TRIzol for gene expression analyses. For immunofluorescence of MECP2 subcellular localization following infection, cells were treated as above, except that cells were grown on coverslips prior to infection and fixation using 4% PFA in PBS. Image acquisition was performed using an apotome microscope (Zeiss).

For assessment of HSV-1 infection by plaque formation assay, 1×10^4^ cells of control of Mecp2-targeting gRNA expressing MEFs were seeded per well of 96-well plates. Twenty-four hours after plating, cells were infected with HSV-1 KOS-64-GFP at MOI 1 in infection medium (DMEM supplemented with 1% FBS, 1% penicillin/streptomycin and 1% Glutamine) for 90 min before medium replacement with DMEM supplemented with 1% human serum, 1% penicillin/streptomycin and 1% Glutamine. Sixteen hours later, medium was replaced with DMEM supplemented with 10% FBS medium,1% penicillin/streptomycin and 1% Glutamine for an additional 32 hours. Cells were either fixed with 4% paraformaldehyde (PFA) prior to staining with crystal violet and plaque counting or supernatants collected to infect WT-MEFs and assessment of infected cells by GFP.

### VSV-GFP infections, quantifications and imaging

VSV-GFP was a gift from Dr Sebastien Pfeffer (IBMC, France). VSV-GFP was amplified on BHK21 cells, aliquoted and frozen at −80°C. Single-round titration of viral stock was performed on NIH-3T3 cells in 96 well plates in technical triplicates. Briefly, cells were infected with serial dilutions of VSV-GFP for 1 h in DMEM 2% FBS, washed with PBS and media replaced. Cells were fixed using 4% PFA for 20 mins at room temperature, permeabilized in 0.2% Triton X-100 for 10 minutes and nuclei stained with DAPI. The percentage of infected cells (i.e. GFP-positive cells) was determined using an ImageXpress Pico (Molecular Devices) and the number of infectious particles per ml was determined. To determine infectivity of VSV-GFP in RAW cells expressing control or Mecp2-targeting gRNAs, the same protocol as above for the NIH-3T3 titration was used.

For confocal imaging experiments, RAW CTRL or Mecp2 KO were plated in either 96 well plates (6×10^5^ cells) or in 24 well plates on glass coverslips (4×10^6^ cells) and infected the next day with the indicated MOIs of VSV-GFP for 7h30 and fixed as above. Cells were permeabilized as above, blocked with 10% Normal Goat Serum (Invitrogen) for 1 h and immunofluorescence was performed. For cells in 96 well plates the cells were treated as above. For cells on glass coverslips, cells were incubated with primary antibodies (Rabbit anti-MECP2, Cell Signaling Technologies, D4F3, 1/50 and Mouse anti-VSVG, Kerafast, Kf-Ab01401-2.0, 1/250) in PBS 1X 0.5% BSA for 1 h followed by incubation with Alexa Fluor secondary antibodies (Thermo Fisher Scientific) and DAPI for 1 h. Cells were washed in water and mounted on slides using ProLong Gold Antifade Mountant (Thermo Fisher Scientific). Imaging was performed on an LSM980 8Y confocal microscope (Zeiss), using a 63x lens for glass coverslips and 10x lens for 96 well plates. Post-processing was performed using the FIJI software ^33^.

### RNA-Seq data analysis

Raw reads were subjected to quality control and adapter trimming using Trimgalore ^34^ (version 0.6.6). Trimmed reads were then aligned to the reference genome (mm9) using HISAT2 (version 2.2.1) ^35^ Converting sam to bam, sorting and indexing bam files have been performed using samtools ^36^ (version 1.11). Aligned sequencing reads have been count using HTSeq tool ^37^ (version 0.11.3; options: -i gene_id --additional-attr gene_name -t exon). Differentially expressed genes analysis have been detected using DESeq2 R package (version 1.32.0) ^38^ Bigwig files have been generated using bamCoverage tool from deepTools suite ^39^ (version 3.4.2; options: -- smoothLength 15 --normalizeUsing RPKM). For transposable element expression (retrotranscriptome) analysis, trimmed reads were mapped to the reference genome (mm9) genome using HISAT2 (version 2.2.1) ^35^. The aligned reads were converted from SAM to BAM format, sorted, and indexed using Samtools (version 1.11). Aligned reads were then counted using Telescope (version 1.0.3.1) ^40^.

### MinION sequencing data

DNA extraction was performed as described in ^41^.

Nanopore sequencing (Oxford Nanopore Technologies) was performed using a Ligation sequencing kit V14 (SQK-LSK114) and the Native barcoding kit 24 (OXNTSQK-NBD114). The sequencing library was loaded onto a Flow cell R10.4.01 (FLO-MIN114) on a MinION Mk1B sequencer. MinKNOW (version 24.02.8, ONT) was used for sequence planning and data acquisition, with the super-accurate basecalling mode enabled. Dorado (version 7.3.11) was used as the basecalling algorithm to produce pod5 and fastQ file. The automatic real time division into passed and failed reads by the MinKNOW as a quality check, removing reads with quality scores < 10. The quality checked reads were demultiplexed and trimmed for adapters and primers using Dorado followed by mappings with the reference genome (mm9) using Minimap2 ^42^ (version 2.28) and then convert from SAM to BAM format, sorted, and indexed using Samtools (version 1.20). The aligend reads were then counted using Telescope (version 1.0.3).

### Image quantification and statistical analysis

Quantification of the cytosolic and nuclear intensity of Mecp2 was performed using the Fiji software. Co-localization analysis was conducted utilizing the JACoP plugin and colocalization test within ImageJ (Fiji).

Statistical analyses were performed using GraphPad Prism version 9. Unpaired or paired t-test were performed as indicated in the figure legends. For correlation analysis, Pearson’s correlation coefficient was used for normally distributed data. The number of independent experimental replicates for each experiment is provided in the figure legends. All data are expressed as mean ± standard error of the mean (SEM). N.S.: non-significant. *P < 0.05, **P < 0.01, ***P < 0.001 and ****P < 0.0001.

## Results

### Mecp2 interacts with cytosolic dsDNA

Previous work has shown that several actors of cytosolic dsDNA recognition, including cGAS, are primarily nuclear in absence of cytosolic dsDNA ^8,43^, suggesting that dsDNA binders are capable of interacting with dsDNAs regardless of their subcellular localisation. We thus hypothesized that MECP2 may also interact with cytosolic dsDNA. To test this hypothesis, we performed a series of DNA pulldown experiments.

First, whole cell extracts from wild-type (WT) mouse embryonic fibroblasts (MEFs) were incubated with streptavidin beads alone, or streptavidin beads on which either 80nt-long 5’biotin-bearing ssDNA (b-ssDNA) or dsDNA (b-dsDNA) were immobilized (*in vitro* pulldowns), prior to assement of bound proteins by Western blot (WB) (Fig. 1A, left). Importantly, both DNA probes are capable of inducing the expected cGAS-STING pathway activation as characterized by increased cGas levels, Sting degradation, and phosphorylating activation of Sting and Irf3 (Fig. S1A) and increased expression of *Interferon* β (*Ifn*β) and *Interleukin 6* (*Il6*), as well as IFN response genes such as the *C-X-C motif chemokine ligand 10* (*Cxcl10*), and *2’-5’-oligoadenylate synthetase 1* (*Oas1*) (Fig. S1B). *In vitro* DNA pulldowns performed using these probes showed robust recruitment of cGas and Mecp2 to dsDNA (Fig. 1A, right). Congruent with previous work, Mecp2 was not associated with ssDNA and the binding of cGas with ssDNA was less efficient when compared to that with dsDNA (Fig. 1A, right) ^44^. Thus, Mecp2 selectively interact with immune stimulatory dsDNA but not ssDNA probes.

**Figure 1.**
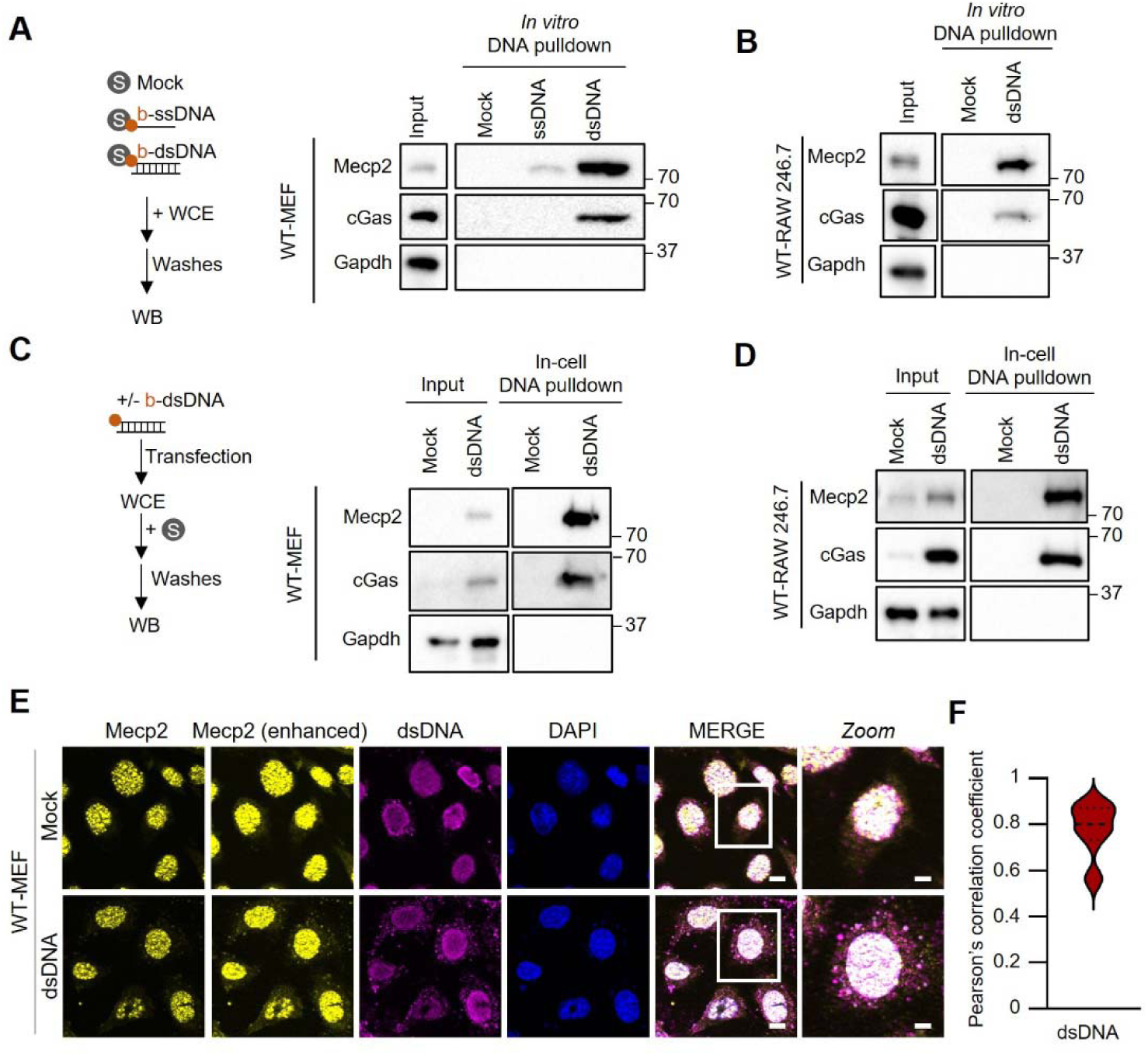
Mecp2 interacts with cytosolic dsDNA. **A** Left: experimental scheme for *in vitro* DNA pull-downs. Right: Whole cell extracts (WCE) prepared from WT-MEF were incubated with streptavidin beads alone or with streptavidin beads-bound biotinylated dsDNA (b-dsDNA) or ssDNA (b-ssDNA) prior to *in vitro* DNA pull-down. Input and eluates were analyzed by Western blot (WB) using the indicated antibodies. **B** As in **A**, except that WCE prepared from WT-RAW264.7 were used. Input and eluates were analyzed by WB using the indicated antibodies. **C** Left: Experimental scheme for in-cell DNA pulldowns. Right: WT-MEF were transfected or not with b-dsDNA before whole-cell extract preparation and pull-down using streptavidin-affinity beads. Input and eluates were analyzed by WB using the indicated antibodies. **D** As in **C**, except that except that WCE were prepared from WT-RAW264.7 cells transfected or not with b-dsDNA. **E** Immunofluorescence analysis was conducted on WT-MEF transfected or not with dsDNA for 6 hours using anti-Mecp2 antibody, anti-dsDNA antibody and DAPI nuclear staining. BF, bright field. Images are representative of 3 independent experiments. Scale bar: 10 µm, except for Zoom: 5µm. **F** Pearson’s correlation coefficient was calculated on the cytosolic dsDNA and Mecp2 signals in dsDNA-stimulated WT-MEF as in **E**. WB and images are representative of at least 3 independent experiments.

We next wished to verify if the interaction of Mecp2 with dsDNA can be recapitulated in other cell types. To this aim, we performed *in vitro* DNA pulldowns using whole cell extracts from the WT-RAW246.7 murine myeloid cell line. Both Mecp2 and cGas were recovered as dsDNA binders in this assay (Fig. 1B). This suggests that the ability of Mecp2 to interact with immune-stimulatory dsDNA is a conserved mechanisms across cell types.

Next, to assess whether recruitment of Mecp2 to dsDNA occurs in cells, MEFs were transfected or not with b-dsDNA for 6 hours, a time point at which immunofluorescence analyses show that transfected dsDNA probes are prominently cytosolic (Fig. S1C). Whole cell extracts were prepared and used in pulldowns using streptavidin affinity beads (in-cell pulldown, Fig. 1C, left). WB analysis of pulled-down material showed that Mecp2 and cGas k,l interact with dsDNA in cells (Fig. 1C, right). Similar results were obtained in WT-RAW264.7 (Fig. 1D) and in the THP-1 human myeloid cell line (Fig. S1D) when in-cell pulldowns were performed 6 hours after transfection of b-dsDNA. These data thus show that Mecp2 can interact dsDNA in MEFs as well as in immune murine and human cell lines. These data further suggest that the interaction takes place in the cytosol.

To further assess where the interaction between Mecp2 and dsDNA probes takes place, we performed *in vitro* DNA pulldowns using cytosolic and nuclear extracts from MEFs (Fig. S1E, left). WB analyses showed an enrichment of Mecp2 bound to dsDNA in both the cytosolic fractions (Fig. S1E, right), supporting and interaction of cytosolic Mecp2 with dsDNA. In addition, we performed immunofluorescence assays where WT-MEFs were challenged with dsDNA prior to staining using Mecp2 and dsDNA-specific antibodies. We observed that dsDNA challenge led to Mecp2-specific staining in the cytosol (Fig. 1E and S1F), with a significant overlap with the dsDNA signal (Fig. 1F). Thus, our data support that Mecp2 interacts with cytosolic dsDNAs.

### Mecp2 is actively exported from the nucleus in the presence of cytosolic dsDNA

Mecp2 is mostly known for its nuclear localization and functions (PMID: 30157418). Thus, its presence in the cytosol is surprising. Quantification of Mecp2 staining showed that dsDNA challenge led to increased cytosolic staining, coupled to decreased nuclear staining (Fig. S1F). This led us to question whether dsDNA stimulation may lead to Mecp2 export from the nucleus.

To test whether increased cytosolic levels of Mecp2 results from active export from the nuclear compartment, we first performed subcellular fractionation experiments where the cytosolic and nuclear soluble fractions were isolated following challenge with dsDNA. WB analyses showed an enrichment of Mecp2 in cytosolic fractions of both WT-MEF (Fig. 2A) and WT-RAW264.7 (Fig. S2A), upon dsDNA challenge. Next, we performed immunofluorescence analyses and assessed Mecp2 subcellular localization in MEF and RAW264.7 following dsDNA challenge. This showed that dsDNA stimulation led to Mecp2 cytosolic staining in both cell types (Fig. 2B top panels, 2C and S2B). In addition, we treated WT-MEFs with the Leptomycin B nuclear export inhibitor prior to stimulation or not with dsDNA. Leptomycin B treatment led to an efficient block of nuclear-cytosolic transport, as shown by immunofluorescence of the RAN Binding Protein 1 (Ranbp1) (Fig. S2C). As expected, Leptomycin B treatment also abolished dsDNA-induced cGas export (Fig. S2D)^43^. We found that Leptomycin B treatment reduced the levels of Mecp2 in the cytosol following dsDNA stimulation (Fig. 2B, bottom panels and 2C). Conversely, Leptomycin B treatment led to increased retention of Mecp2 in the nucleus following dsDNA challenge (Fig. 2B, bottom panels and 2C). These data support that cytosolic Mecp2 accumulation upon dsDNA challenge is driven by active export of Mecp2.

**Figure. 2.**
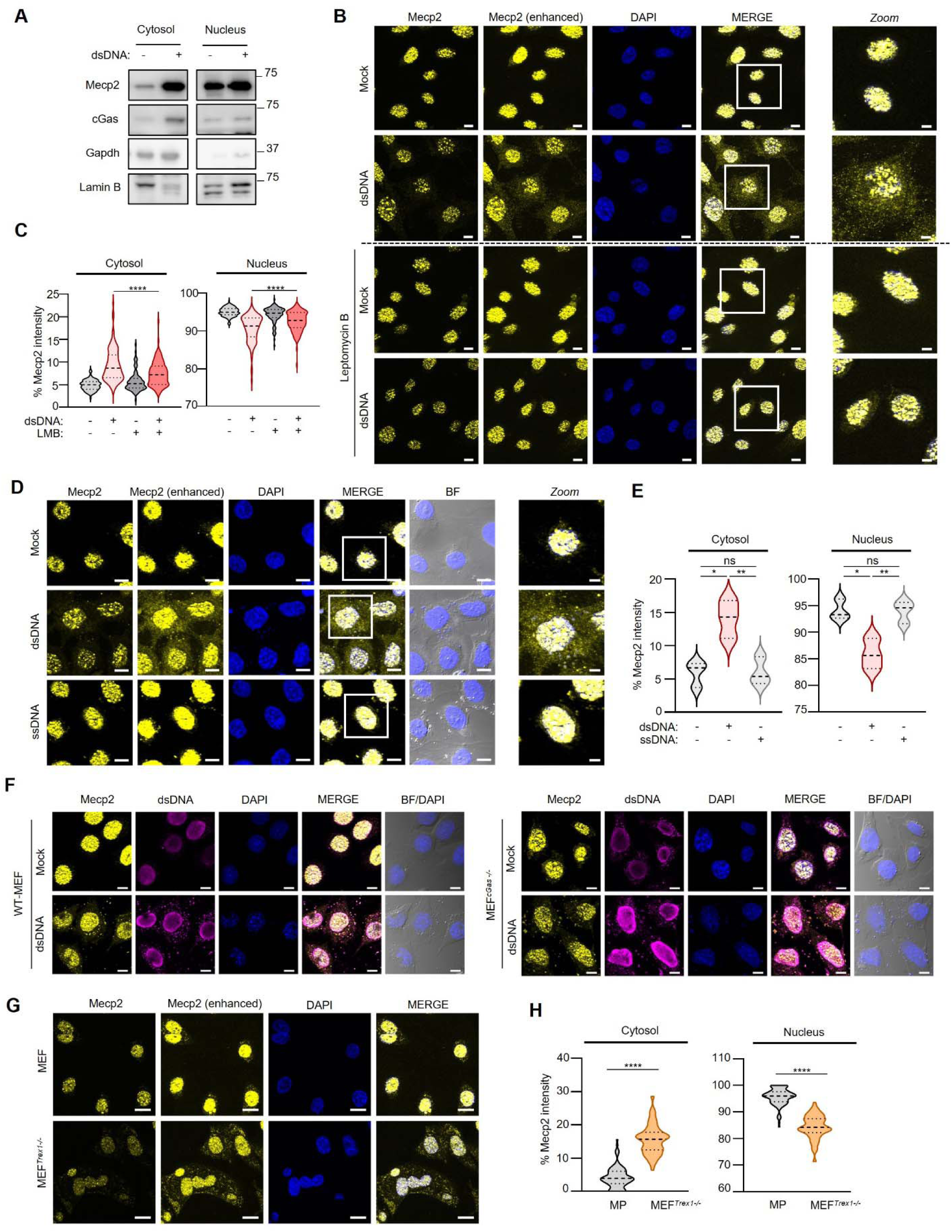
Challenge with dsDNA triggers Mecp2 export. **A** Cytosolic and nuclear soluble fractions were prepared from WT-MEF transfected or not with dsDNA for 6 hours. Fractions were analyzed by WB using indicated antibodies. WB is representative of 3 independent experiments. **B** Immunofluorescence analysis was performed on WT-MEF treated or not with 20 nM of Leptomycin B (LMB) prior to dsDNA transfection for 3 hours using anti-Mecp2 antibody (enhanced signal or not) and DAPI nuclear staining. Scale bar: 10µm; Scale bar for Zoomed images: 5µm. Images are representative of two independent experiments. **C** Violin plots show the % of Mecp2 intensity in the cytosol in cells treated as in **B**; n=95 cells per condition. **D** Immunofluorescence analysis of WT-MEF transfected or not with dsDNA and ssDNA for 6 hours was performed using anti-Mecp2 antibody (enhanced signal or not) and DAPI nuclear staining. BF, bright field. Scale bar: 10µm; Scale bar for Zoomed images: 5µm. Images are representative of 3 independent experiments. **E** Violin plots show the % of Mecp2 intensity in the cytosol and in the nucleus in experiments performed as in **D**; n=105 cells per condition. **F** Immunofluorescence analysis was conducted on WT-MEF and MEF*^cGas-/-^* cells transfected or not with dsDNA for 6 hours using anti-Mecp2 antibody, anti-dsDNA antibody and DAPI nuclear staining. BF, bright field. Images are representative of 3 independent experiments. Scale bar: 20 µm. **G** Immunofluorescence analysis was performed on WT-MEF and MEF*^Trex1-/-^* using anti-Mecp2 antibody and DAPI nuclear staining. Scale bare: 20 µm. Images are representative of two independent experiments. **H** Violin plots show the % of Mecp2 intensity in the cytosol and nucleus of cells treated as in **a**, n=97 cells per condition. Significance was assessed using Student T-test. ns: non-significant. *P < 0.05, **P < 0.01, ***P < 0.001 and ****P < 0.0001.

To assess whether Mecp2 export is specific to dsDNA as suggested by Fig. 1A, we next challenged WT-MEF with ssDNA or dsDNA prior to analysis of Mecp2 subcellular localization. Immunofluorescence analyses showed an increase of Mecp2 cytosolic levels, coupled to a decrease of Mecp2 nuclear levels only upon challenge with dsDNA, and not with ssDNA (Fig 2D-E). This supports that Mecp2 export from the nucleus is a process triggered by the presence of cytosolic dsDNA.

Both dsDNA or ssDNA challenge are contexts where inflammatory signalling are involved (Fig S1A-B). That Mecp2 is not displaced in the cytosol following ssDNA stimulation suggests that inflammatory signalling does not drive Mecp2 export. To confirm this observation, we assessed Mecp2 subcellular localization in a context where dsDNA-associated inflammatory responses are abrogated. We thus challenged WT-MEF and cGas-knockout MEFs (MEF*^cGas-/-^*) with dsDNA. Immunofluorescence analysis of Mecp2 subcellular localization in response to dsDNA stimulation showed that Mecp2 is exported to the cytosol regardless of cGas expression (Fig. 2F). These data thus indicate that Mecp2 export is trigger by the presence of dsDNA in the cytosol and is not a response to cGas-Sting signalling.

Finally, we questioned whether cytosolic accumulation of endogenous DNA is sufficient to trigger Mecp2 export. To this aim, we performed Mecp2 subcellular localization analysis in MEFs harboring an invalidating mutation in the *Three prime repair exonuclease 1* (Trex1) gene. Indeed, Trex1 is a nuclease that degrades cytosolic DNAs. Invalidating mutations in Trex1 lead to chronic accumulation of cytosolic DNAs ^45^, notably owing to aberrant activity of the long interspersed element-1 (LINE-1) endogenous retroelement. Immunofluorescence analysis of control and Trex1-deficient MEFs showed the presence of Mecp2 in the cytosol (Fig. 2G-H). Furthermore, treatment with the Tenofovir reverse transcriptase inhibitor, known to inhibit LINE-1 activity ^46^, led to decreased cytosolic dsDNA, cGas (Fig. S2E) and Mecp2 staining (Fig. S2F-G). Therefore, our data show that the presence of endogenous cytosolic dsDNA is sufficient to trigger Mecp2 export. Moreover, these data show that this process can be reversed by inhibition of reverse transcriptase activities.

### Mecp2 and cGas interact with the same dsDNA moieties in the cytosol

Experiments performed in Fig.1 and S1 show that Mecp2 and cGas are capable of binding dsDNA probes *in vitro* and in cells. We thus questioned whether they may bind the same dsDNA species in the cytosol.

To this aim, MEFs engineered to stably express an enhanced green fluorescence protein (EGFP)- tagged cGas allele (MEF*^EGFP-cGas^*) were transfected or not with dsDNA prior to immunofluorescence analyses of Mecp2 and EGFP-cGas co-localization using Pearson’s correlation coefficient. This showed that dsDNA transfection increased the colocalization between cGas and Mecp2 in the cytosol (Fig. 3A), suggesting tripartite interaction between dsDNA, cGas and Mecp2.

**Figure 3.**
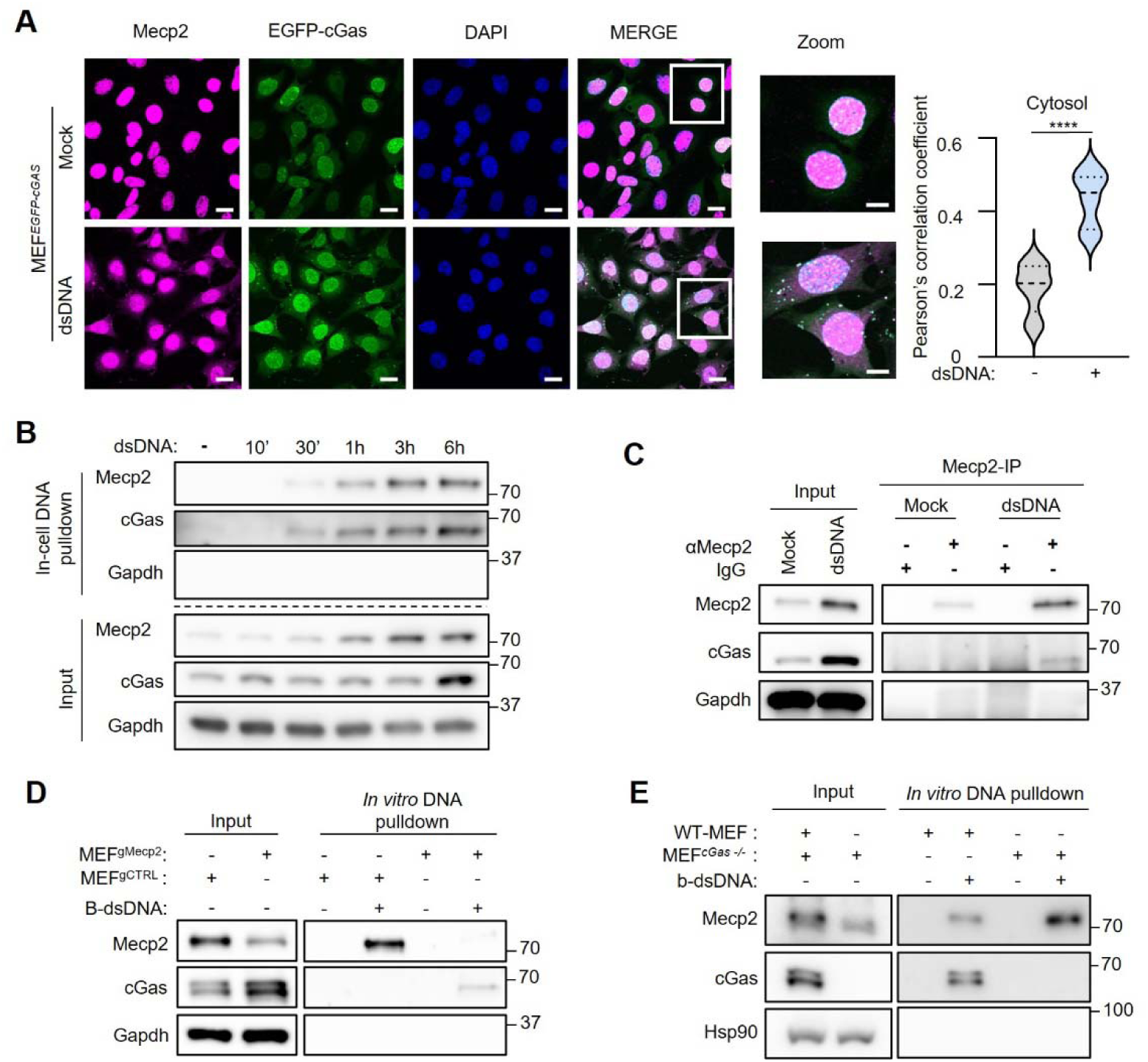
Absence of Mecp2 incrases cGAS interaction with cytosolic dsDNA. **A** Left: Immunofluorescence analysis was conducted on MEF stably expressing a GFP-cGAS construct (MEF*^GFP-cGAS^*) transfected or not with dsDNA for 6 hours using anti-Mecp2 antibody and DAPI nuclear staining.. Images are representative of 3 independent experiments. Scale bar: 20 µm; Scale bar for Zoom: 10 µm. (Right) Graph presents mean ± SEM Pearson’s correlation coefficient values for co-localization of cGas and Mecp2. p values were determined by Student’s t test. ****p < 0.0001; n=12. **B** WT-MEF were transfected or not for 10 min, 30 min, 1 hour, 3 hour or 6 hours with b-dsDNA before whole-cell extract preparation and pull-down using streptavidin-affinity beads. Input and eluates were analyzed by WB using the indicated antibodies. **C** WT-MEF were transfected or not with dsDNA before whole-cell extract preparation and immunoprecipitation using control IgG or a Mecp2-specific antibody. Input and immunoprecipitated material were analyzed by WB using the indicated antibodies. **D** Whole cell extracts prepared from WT-MEF or MEF*^cGas-/-^* cells were incubated with streptavidin beads alone or with streptavidin bead-bound b-dsDNA prior to pull-down. Input and eluates were analyzed by WB using the indicated antibodies. **E** Whole cell extracts prepared from MEF expressing a control non-targeting gRNA (MEF^gCTRL^) or a Mecp2-targeting gRNA (MEF^gMecp2^) were incubated with streptavidin beads alone or with streptavidin bead-bound b-dsDNA prior to pull-down. Input and eluates were analyzed by WB using the indicated antibodies. WB is representative of 3 independent experiments.

We next interrogated the dynamics of the interaction of Mecp2 and cGas with cytosolic dsDNA. To this aim, WT-MEFs were transfected with b-dsDNA for 10 minutes, 30 minutes, 1 hour, 3 hours and 6 hours prior to whole cell extraction and in-cell DNA pulldowns using streptavidin affinity beads. WB analyses showed Mecp2 and cGas interaction with dsDNA as early as 30 min post transfection (Fig. 3B). This suggests similar interaction dynamics of Mecp2 and cGas with dsDNA. This was further tested by performing Mecp2 immunoprecipitation in WT-MEFs transfected or not with dsDNA prior to assessment of cGas co-immunoprecipitation. We found that dsDNA transfection led to co-immunoprecipitation of cGas with Mecp2, as compared to mock transfection (Fig. 3C). Thus, when taken together, these data support that the presence of cytosolic dsDNA triggers the formation of Mecp2, cGas and dsDNA containing complexes.

The recruitment of cGas and Mecp2 to the same dsDNA molecules in the cytosol raises the possibility that they may compete for interaction. To test this, we first used WT-MEF and MEF*^cGas-/-^* to perform *in vitro* DNA pulldowns. WB analyses showed that absence of cGas enhanced the recruitment of Mecp2 to dsDNA (Fig. 3D and Fig. S3). Conversely, we assessed whether Mecp2 regulates cGas recruitment to dsDNA. To this aim, we used MEFs expressing control (MEF^gCTRL^) or Mecp2-targeting gRNAs (MEF^gMecp2^) to perform *in vitro* dsDNA pulldowns. We found that reduced levels of Mecp2 led to increased cGas recruitment to dsDNA (Fig. 3E). This suggests that cGas and Mecp2 bear the capacity to regulate their respective recruitment to cytosolic dsDNA, further supporting that they are recruited to the same dsDNA molecules.

### Mecp2 inhibits cGas-dependent Type I Interferon responses

Our data show that the absence of Mecp2 leads to increase of cGas recruitment to cytosolic dsDNA (Fig. 3), suggesting that absence of Mecp2 may promote enhanced dsDNA-induced cGas-dependent signalling. While it was previously reported that absence of Mecp2 led to increased inflammatory responses, a role of Mecp2 in regulating cGas- or dsDNA-associated inflammatory responses was not reported.

To assess whether Mecp2 may regulate cGas activity, we challenged MEF^gCTRL^ and MEF^gMecp2^ with dsDNA prior to assessment of cGas-dependent pathway activation. Immunofluorescence analyses of cGas subcellular localization in these cell lines showed that dsDNA stimulation enhanced cGas levels in the cytosol in MEF^gMecp2^as compared to MEF^gCTRL^ (Fig. 4A). Assessment of intracellular cGAMP levels in MEF^gCTRL^ and MEF^gMecp2^ cell lines showed that low levels of Mecp2 led to higher levels of dsDNA-associated 2’3’-cGAMP levels (Fig. 4B). Analyses of the expression of genes classically associated with cGas activation, such as *Ifn*β, *Il6* and *Cxcl10* showed that Mecp2 knockout led to their increased expression (Fig. 4C). Similar data were obtained in RAW264.7, where Mecp2 knockout (RAW264.7^gMecp2^) led to enhanced dsDNA-induced expression of *Ifn*β, and *Interferon-stimulated gene 15* (*Isg15*) as compared to control cells (RAW264.7^gCTRL^) (Fig. S4), supporting the conservation of an inhibitory role of Mecp2 in different cell lines. Importantly, the inflammatory responses witnessed upon dsDNA stimulation in cells expressing a Mecp2-targeting gRNA is decreased by treatment with the H-151 Sting inhibitor (Fig. 4D), supporting that the witnessed inflammatory signature is, at least in part, Sting-dependent. Finally, we transfected MEFs with either control (Empty) or Mecp2-expressing vector (eMecp2). We found that overexpression of Mecp2 prior to dsDNA stimulation led to decreased expression of *Ifn*β and *Il6*, as compared to Empty vector-expressing cells (Fig. 4E). This confirms the role of Mecp2 as a negative regulator of cGas-associated inflammatory responses.

**Figure 4.**
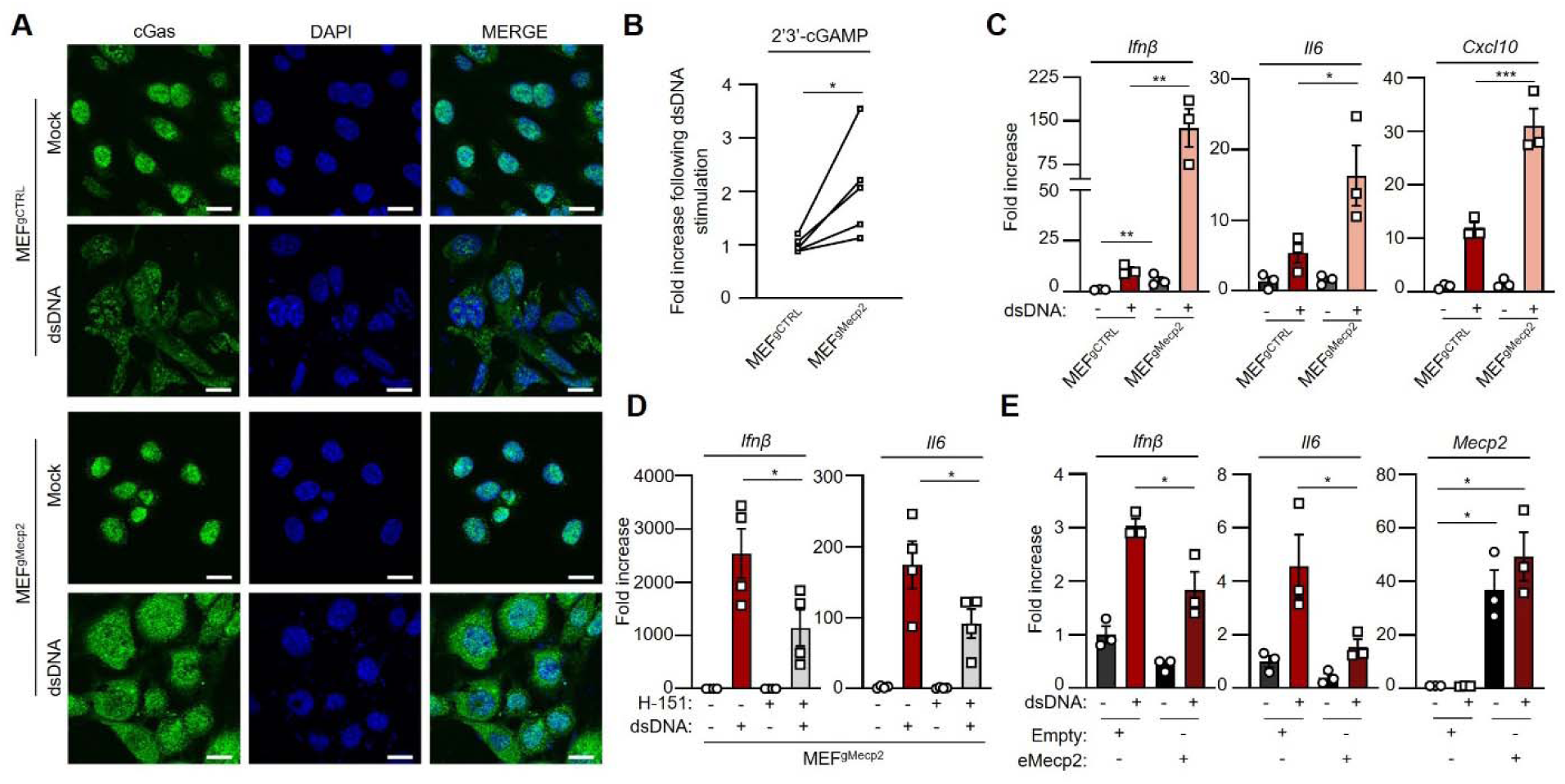
Absence of Mecp2 primes cGAS-depednent Inflammatory responses. **A** Immunofluorescence analysis was conducted on MEF^gCTRL^ or MEF^gMecp2^ after challenged or not with dsDNA for 6 hours, using anti-cGas antibody and DAPI nuclear staining. Scale bar: 20 µm. Images are representative of 3 independent experiments. **B** Intracellular cGAMP levels were measured by ELISA following transfection or not with dsDNA of MEF^gCTRL^ or MEF^gMecp2^. Graph presents fold increase cGAMP levels from 5 independent experiments. **C** MEF^gCTRL^ or MEF^gMecp2^ were challenged or not with dsDNA for 6 hours prior to gene expression analysis. Graph presents mean (± SEM) *Ifn*β, *Cxcl10* and *Il6* mRNA levels (n=3 independent experiments). **D** MEF^gMecp2^ were challenged or not with dsDNA for 6 hours in the presence or not of the H-151 Sting inhibitor. Graphs present mean (±SEM) *Ifn*β, and *Il6* mRNA levels (n=4 independent experiments). **E** MEF overexpressing (eMecp2) or not (Empty) WT-Mecp2 were transfected or not with dsDNA for 6 hours prior to analysis of *Ifn*β, *Il6* and *Mecp2* mRNA levels. Graphs present mean (±SEM) from 3 independent experiments. Significance was assessed using Student T-test. ns: non-significant. *P < 0.05, **P < 0.01, and ***P < 0.001.

Taken together, these data (Fig. 3 and 4) show that the absence of Mecp2 leads to increased cGas levels, which results in increased cGas recruitment to cytosolic dsDNA in cells and subsequent activation of cGas-dependent inflammatory responses.

### Absence of Mecp2 enforces an antiviral state

That Mecp2 inhibits dsDNA-associated inflammatory responses suggests that absence of Mecp2 may lead to chronic low-grade type I IFN responses. The latter are known to lead to the establishment of protective antiviral states. This led us to question the interplay between viral infections and Mecp2 (Fig. 5A).

**Figure 5.**
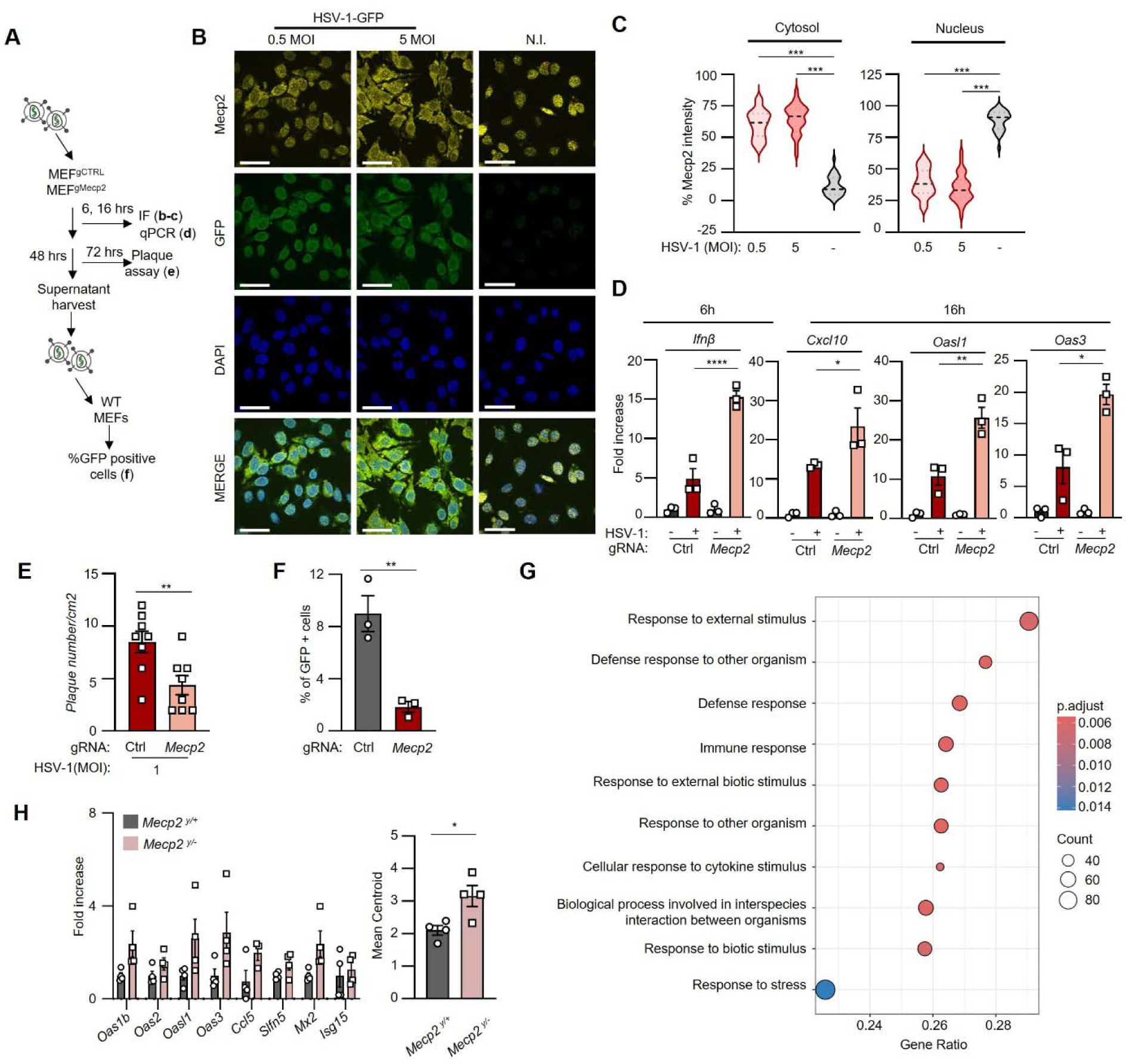
Absence of Mecp2 enforces an antiviral state. **A.** Experimetna scueme for **B-F. B** WT-MEF cells infected or not with HSV-1-GFP at 0.5 or 5 multiplicity of infection (MOI) were subjected to immunofluorescence using anti-Mecp2 antibody and DAPI nuclear staining. Images are representative of 3 independent experiments. Scale bar: 50µm. **C** Violin plots show the % of Mecp2 intensity in the cytosol in cells infected as in **B**; n=95 cells per condition. **D** MEF^gCTRL^ or MEF^gMecp2^ were infected or with 5 MOI of HSV-1-GFP for 6 and 16 hours prior to gene expression analysis. Graphs present the mean ± SEM *Ifn*β, *Cxcl10*, *Il6*, *Oasl1* or *Oas3* mRNA levels (n= 3 independent experiments). **E** Plaque numbers were counted following 72h of infection with 1 MOI of HSV-1-GFP of MEF^gCTRL^ or MEF^gMecp2^. Graph presents mean (±SEM) plaque number per cm2 as calculated in 8 replicates, representative of 3 independent experiments. **F** Graph presents the mean (±SEM) percentage of GFP positive cells measure in cells treated as in **E** (n=3 independent experiments). **G** Gene Set Enrichment Analysis (GSEA) was performed using clusterProfiler (version 3.8) looking for Biological Process (BP) on DESeq2 results (log2foldchange>0.01. Gsea pvalueCutoff = 0.05 with scoreType = pos). **H** Gene expression analysis was performed in livers of male Mecp2^+/y^ and Mecp2^-/y^ mice. Left: Graph present mean (±SEM) fold increase gene expression in Mecp2^-/y^ mice as compared to Mecp2^+/y^; n=4 mice per group. Right: mean centroid expression was calculated based on the expression of indicated genes (n=7). Significance was assessed using Student T-test. ns: non-significant. *P < 0.05, **P < 0.01, ***P < 0.001 and ****P < 0.0001.

We first questioned whether the delivery of DNA in the cytosol through infection with a DNA virus would induce accumulation of Mecp2 in the cytosol as witnessed upon dsDNA transfection (Fig. 2). We used a molecular clone of the KOS64 Herpes simplex virus 1 (HSV-1), harbouring a dsDNA genome, expressing a GFP reporter (HSV-1-GFP). We found that infection of WT-MEFs with HSV-1-GFP led to the presence of Mecp2 in the cytosol (Fig. 5B-C). Similar experiments were conducted using a molecular clone of the Vesicular stomatitis virus (VSV), a virus harbouring a ssRNA genome, expressing a GFP reporter (VSV-GFP). In contrast to what was visualized following infection with HSV-1-GFP, we found that infection with VSV-GFP did not lead to cytosolic staining of Mecp2 (Fig. S5A). These experiments show that the delivery of dsDNA in the cytosol is necessary and sufficient to induce Mecp2 nuclear export.

Next, we assessed whether the presence or absence of Mecp2 can influence inflammatory responses following infection with HSV-1-GFP. To this aim, MEF^gCTRL^ and MEF^gMecp2^ were infected with HSV-1-GFP for 6 and 16 hours prior to assessment of the expression of *Ifn*β at 6 hours and of antiviral IFN-stimulated genes such as *Cxcl10*, as 2’-5’-Oligoadenylate Synthetase-Like (*Oasl1*) and 2’-5’-oligoadenylate synthetase 3 (*Oas3*) at 16 hours post infection. Absence of Mecp2 led to increased expression of those genes following HSV-1-GFP infection (Fig. 5D). Thus, lower levels of Mecp2 leads to increased expression of inflammatory genes following infection with HSV-1-GFP, confirming the inhibitory impact of Mecp2 on cGas-associated signalling (Fig. 4).

We next assessed the capacity of HSV-1 to infect and replicate in Mecp2-deficient and proficient cells. To this aim, MEF^gCTRL^ and MEF^gMecp2^ were infected with HSV-1-GFP for 72 hours prior to assessment of plaque formation. Decreased levels of Mecp2 led to decreased plaque formation (Fig. 5E and S5B), attesting to reduced ability of HSV-1-GFP to infect these cells as compared to control cells. Next, HSV-1-GFP produced from MEF^gCTRL^ and MEF^gMecp2^ were used to infect WT-MEF cells prior to quantification of GFP-positive cells. We found that infection of WT-MEF with the supernatant collected from MEF^gMecp2^ led to decreased number of GFP-positive cells as compared to supernatant collected from MEF^gCTRL^ (Fig. 5F and S5C). In contrast, when similar experiments were conducted using VSV-GFP, we found that absence of Mecp2 did not significantly alter viral infection (Fig. S5D). Altogether, these data show that absence of Mecp2 fosters an antiviral state that hinders infection by HSV-1. This further suggests that absence of Mecp2 is sufficient to foster an antiviral state that is efficient towards DNA virus infection.

Since the presence of a type I IFN antiviral signature was not previously reported in Mecp2 deficiency, we finally assessed whether Mecp2 knockout is sufficient to promote the expression of a type I IFN signature. We first performed RNA sequencing (RNAseq) on RAW264.7^gCTRL^ and RAW264.7^gMecp2^. Gene Set Enrichment Analysis (GSEA) was conducted on genes upregulated in RAW264.7^gMecp2^ as compared to RAW264.7^gCTRL^, revealing an upregulation of processes related to stress response, immune response and biological interactions between organisms, including responses to external and biotic stimuli (Fig. 5G). We next assessed the presence of such an antiviral signature in the liver of *in vivo* mouse models of Mecp2 deficiency (Mecp2^y/-^) as compared to WT littermates (Mecp2^y/+^). Analysis of the expression levels of type I IFN responses genes and antiviral genes (*Oas1b*, *Oas2*, *Oasl2*, *Oas3*, *C-C motif chemokine ligand 5* (*Ccl5*), *Schlafen 5* (*Slfn5*), MX dynamin-like GTPase 2 (*Mx2*) and *Isg15*) showed a tendency for increased expression (Fig. 5H, Left). Mean centroid analysis of those antiviral genes showed a significant increase in Mecp2 deficient animals (Fig. 5H, Right), confirming the presence of an antiviral type I IFN response in Mecp2 deficiency.

Thus, our data suggest that absence of Mecp2 therefore leads to enhanced cGas activity in the presence of cytosolic dsDNA, enforcing an antiviral state affecting dsDNA virus infection. These data thus further support that Mecp2 is a negative regulator of cGAS activity.

## Displacement of Mecp2 from the nucleus disrupts its canonical function

We finally questioned whether Mecp2 nuclear export following dsDNA stimulation may disrupt its canonical gene expression regulation function. Owing to the abundance of Mecp2, the presence of multiple targets in the genome, and differential methylation patterns between cell types, identification of the precise regions regulated by MeCP2 has proven difficult. Genes consensually identified as repressed by Mecp2 notably comprise genomic locations corresponding to endogenous retroelements ^28,47,48^. We thus decided to assess LINE-1 expression following dsDNA stimulation to determine whether Mecp2 canonical function is altered.

To this aim, RAW264.7 cells were transfected with dsDNA prior to RNAseq. The retrotranscriptome was analyzed using the Telescope software tool ^28^ in order to identify reads corresponding to endogenous retroelements. Principal component analysis was conducted on all expressed genes and on the retrotranscriptome, showing that Mock and dsDNA stimulated samples segregate into distinct subpopulations (Fig. S6A-B). Volcano plots showed that alongside the expected upregulation of gene expression following dsDNA stimulation (Fig. S6C), dsDNA stimulation led to a robust upregulation of sequences corresponding to endogenous retroelements (726 unique transcripts; Fig. 6A). Obtained reads were normalized to their relative abundance in the genome prior to analysis of the global change in endogenous retroelement gene expression. This revealed a significant increase in reads corresponding to class I retroelements (Fig. 6B), which are those which replicate through RNA intermediates ^29^. Significantly upregulated class I retroelements include long terminal repeats (LTR), endogenous retroelements (ERV) and non-LTR transposons. Non-LTR transposons significantly increased upon dsDNA stimulation comprise LINEs, but also Short interspersed nuclear elements (SINEs) (Fig. 6B). We next examined more closely the upregulation of LINE-1 upon dsDNA stimulation and found that there is a significant increase of LINE-1 expression upon dsDNA stimulation as compared to the mock condition (Fig. 6C).

**Figure 6.**
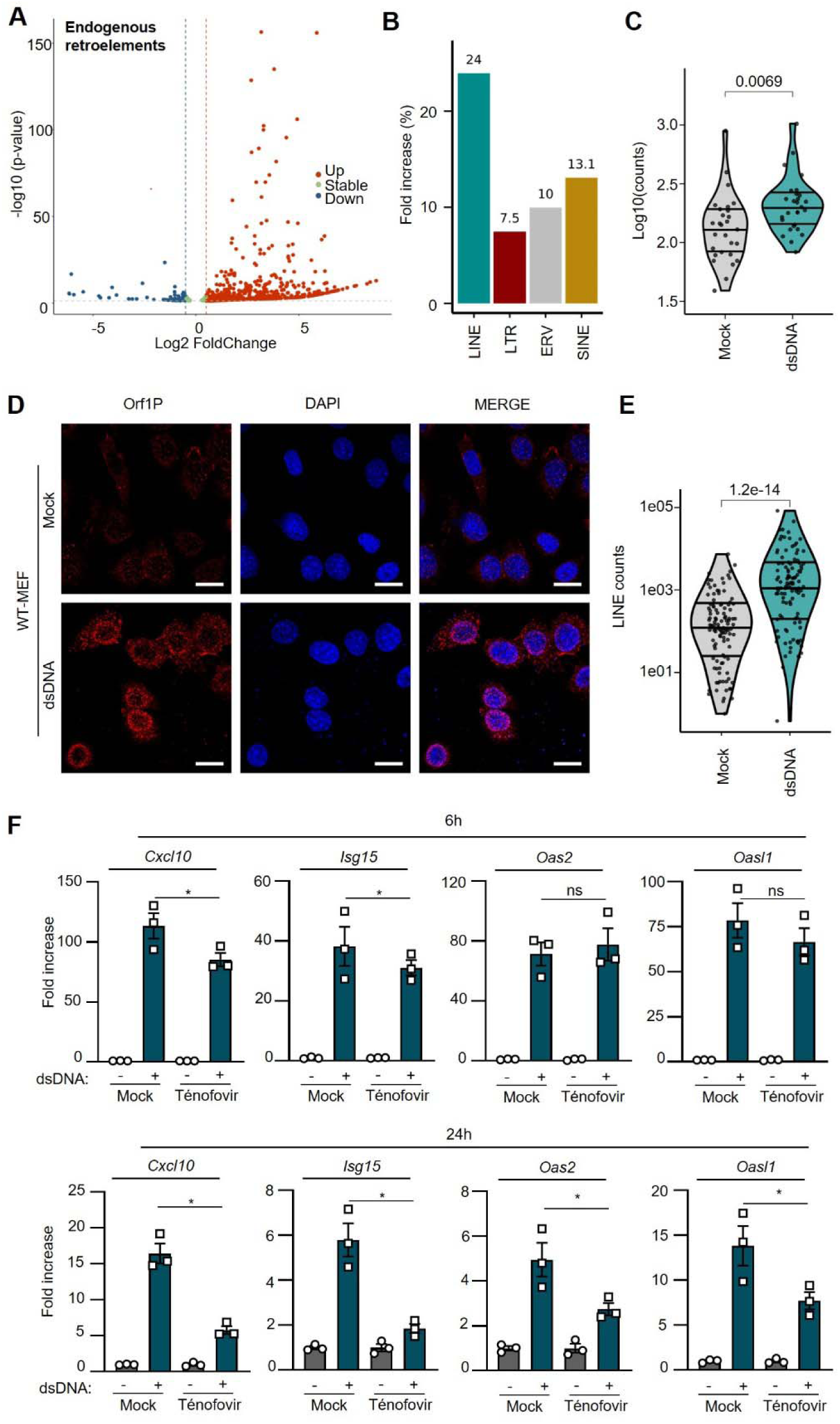
Mecp2 deficiency leads to accumulation of immunogenic LINE-1 derived DNA. A Volcano plot representing upregulated, stable or downregulated transcripts corresponding to endogenous retroelements WT-RAW264.7 stimulated with dsDNA for 6hours as compared to non-stimulated cells. **B** Graph presents the % fold increase in trascripts corresponding to type I endogenous retroelements in data from **A**. LINE: long interspersed nuclear elements, LTR: Long terminal repeats containing retroelements; ERV: Endogenous RetroViruses; SINE: short interspersed repetitive elements. C Violin plots present the transcripts corresponding to LINE-1 elements in data from **A**. **D** Immunofluorescence analysis of WT-MEF transfected or not with dsDNA for 6 hours using anti-Orf1p antibody and DAPI nuclear staining. Scale bar: 20µm. Images are representative of 3 independent experiments. **E** Cytosolic DNA isolated from WT-MEF treated as in **D** were sequenced using nanopore sequencing. The violin plot presents the reads correspondinf to LINE-1. **F.** WT-MEF were transfected or not with dsDNA for 6 or 24 hounrs in the presence of the tenofovir reverse transcriptase inhinbitor. The graphs present the mean fold increase of *Cxcl10*, *Isg15*, *Oas2* and *Oasl1* as compared to untreated cells. Significance was assessed using Student T-test. ns: non-significant. *P < 0.05, **P < 0.01, ***P < 0.001.

While Mecp2 has been reported to bind and repress LINEs, Mecp2 was shown to not directly regulate SINEs ^30^. SINEs are rather mobilised by LINE-1 activity ^31^. The upregulation of SINEs (Fig. 6B) could therefore reflect the increased LINE-1 activity witnessed following dsDNA transfection, further suggesting that functional LINE-1-associated proteins may be produced following dsDNA stimulation. To test this hypothesis, immunofluorescence analyses were conducted to assess the levels of the Orf1p protein, which is an essential component of the LINE-1 retrotransposition machinery ^32^. We found that stimulation with dsDNA was sufficient to lead to increased Orf1p levels in the cytosol of WT-MEFs (Fig. 6D and S6D). Therefore, our data indicate that dsDNA stimulation leads to the expression of endogenous retroelements. Owing to the role of Mecp2 in mediating their repression, this suggests that dsDNA stimulation disrupts Mecp2 canonical function.

### LINE-1 derived dsDNA promotes cGAS-signalling

Interestingly, Orf1p bears reverse transcriptase activity. Therefore, increased Orf1p levels may lead to the accumulation of LINE-1-derived DNA in the cytosol. Such DNA species have been shown to be immune-stimulatory, raising the possibility that dsDNA stimulation may lead to an upregulation of LINE-1-derived DNA species that could sustain pro-inflammatory signaling.

To test this hypothesis, we purified cytosolic DNA from WT-MEF stimulated or not with dsDNA and performed DNA sequencing using the nanopore technology. Analysis of LINE-1 sequences showed a 10-fold increase in the cytosolic fraction of cells following dsDNA stimulation (Fig. 6E). This supports that dsDNA stimulation leads to de-repression of LINE-1 expression coupled to an increase of Orf1p and subsequent accumulation of LINE-1 DNA.

Finally, we tested the immunogenic potential of those DNA species by treating dsDNA-stimulated cells with the Tenofovir reverse transcriptase inhibitor. To this aim WT-MEFs and WT-RAW264.7 cells were transfected or not with dsDNA in the presence or not of Tenofovir. Analysis of the expression of *Cxcl10*, *Isg15*, *Oas2* and *Oasl1* was conducted at 6 and 24 hours post stimulation with dsDNA. This showed that, while Tenofovir treatment had a mild impact on the dsDNA-induced expression of those genes at 6 hours, 24 hours post-stimulation, treatment with Tenofovir dampened their induction (Fig. 6F). These data suggest that LINE-1 associated reverse transcriptase activity is involved in promoting persistent type I IFN responses. This further suggests that Mecp2 displacement from the nuclear compartment and the associated upregulation in LINE-1 activity, serves to promote inflammatory signal persistence.

## Discussion

We here provide evidence for a role of Mecp2 as a negative regulator of dsDNA-induced inflammatory responses. Indeed, we show that the presence of cytosolic dsDNA is sufficient to trigger Mecp2 re-localization to the cytosol where it interacts with dsDNA and regulated cGas-associated signalling, dampening type I IFN responses.

This process modifies the cellular transcriptional landscape and primes the expression of genes that are otherwise repressed by Mecp2, yielding both the establishment of an antiviral state and the expression of transposable elements belonging to the abundant LINE-1 family ^28,47,48^. LINE-1 transposons were previously shown to contribute to RTT onset and were suggested as causative of neurologic symptoms ^49^. Interestingly, uncontrolled LINE-1 activity is known to promote the accumulation of cytosolic DNAs that trigger cGAS-STING-dependent inflammatory responses ^29,45,50^. The de-repression of endogenous retroelements induced by dsDNA challenge that we measure in our experiments is both indicative of disruption of MECP2 canonical function, but also serves to shape inflammatory responses, promoting their persistence. Furthermore, our data indicate that treatment with reverse transcriptase inhibitors that prevent LINE-1 DNA accumulation prevents the persistence of type I IFN responses. Knowing the deleterious impact of chronic type I IFN responses, this suggests that reverse transcriptase activity inhibition may permit resolution of the signalling. This further suggests that treatment with reverse transcriptase inhibitors may alleviate inflammation in RTT and therefore reduce the burden of RTT symptoms.

Additionally, our data reveal aberrant cytosolic Mecp2 subcellular localisation in the Trex1-deficiency model of type I interferonopathy. This suggests that chronic IFN responses witnessed in Trex1-deficiency may be a result of Mecp2 -driven de-repression of LINE-1s, and subsequent activation of inflammatory and antiviral genes. In support of this, previous work has shown that treatment with RT inhibitors efficiently prevented endogenous retroelement activity in this model and decreased inflammatory responses and symptoms *in vivo* ^46^ and in patients ^51^. Whether Mecp2 mislocalization plays a role in these contexts remains to be investigated. Furthermore, recent reports indicate that endogenous retroelement activity may be involved in the regulation of tonic inflammatory responses ^52^, suggesting that the absence of Mecp2 may contribute to priming inflammatory responses. Our study shows that Mecp2 ablation led to the upregulation of antiviral genes capable of limiting replication by the HSV-1 dsDNA virus. That replication of the VSV single-stranded RNA virus is not altered by absence of Mecp2 also reinforces that Mecp2 ablation primes cells for dsDNA-associated type I IFN responses.

Alternatively, this may also suggest that the antiviral status elicited by absence of Mecp2 can be overcome by RNA viruses. Indeed, the current view suggests that ssRNA viruses efficiently overcome or circumvent cGas-Sting activation ^53^. This further supports of our evidence that cGas-Sting activity drives the antiviral status in Mecp2-deficiency.

Finally, we report that the presence of a antiviral type I IFN signature in the liver of *in vivo* models of RTT. Interestingly, metabolic dysfunctions were previously reported in livers of Mecp2-deficient mice ^54^. In light of previous work showing that Sting is a crucial regulator of polyunsaturated fatty acid (PUFA) metabolism ^55^ it can be speculated that metabolic alterations observed in Mecp2-deficient mice livers may be attributed to chronic cGas-Sting activation driving PUFA metabolism imbalances. In this line, it has already been proposed that supplementation in Omega-3 polyunsaturated fatty acids may alleviate symptoms of RTT ^22^. It is therefore tempting to propose that Sting pharmacological inhibition may bear promises for RTT patients.

## Acknowledgments

We thank all members of the Molecular Basis of Inflammation laboratory for their critical reading of this manuscript. We thank Denis Tempe for discussions. We thank Soren Paludan for WT and cGas-deficient MEF cell lines and HSV-1-GFP, J Rewhinkel for Trex1-deficient MEFs, Olivier Moncorgé for BHK21 cells and Sébastien Pfeffer for VSV-GP. We acknowledge the MRI imaging facility, member of the national infrastructure France-BioImaging infrastructure supported by the French National Research Agency (ANR-10-INBS-04, “Investments for the future”). We thank Julian Venables for edition of the manuscript.

This work was co-funded by the European Union (ERC, SENTINEL 101087092 to NL and DELV 101039538 to KM). Views and opinions expressed are however those of the author(s) only and do not necessarily reflect those of the European Union or the European Research Council. Neither the European Union nor the granting authority can be held responsible for them. This work was also co-funded by LA LIGUE pour la recherche contre le cancer [AAPARN 2021.LCC/JuF (NL) and PhD fellowship (HC)], the Agence Nationale de Recherche sur le SIDA et les Hépatites virales (ANRS) [ECTZ117448 (NL)], the AFSR (Association Française du Syndrome du Rett) (EV, ZH), the I-SITE Excellence Program of the University of Montpellier, under the Investissements France 2030 [RETTiNA (NL), ChoiCe (NL) and PhD Fellowship (SG)], the Fondation ARC [ARCPJA2021060003720 COPALYS (NL) and PhD fellowship (HC)], La Région Languedoc Roussillon [Prématuration 2021 MODULON 21015964 (NL)] and the Centre National de La Recherche Scientifique [Prématuration CNRS (NL)].

## Author contributions

Conceptualization: NL; Methodology: HC, IKV, SG, CAS, MS, NL; Investigation: HC, IKV, AC, SG, CAS, MS, MAD, MC, MS*, ZH, JM, PLH; Visualization: HC, NL, MS, CT, PLH; Funding acquisition: NL; Project administration: NL; Supervision: IKV, NL, ES, EV, KM; Writing – original draft: NL; Writing – review & editing: HC, SG, IKV, NL, MS, CAS, ES, MS, EV, JM, KM.

## Materials and correspondence

Correspondence and requests for material should be addressed to Nadine Laguette.

## References

1 Miller, K. N. et al. Cytoplasmic DNA: sources, sensing, and role in aging and disease. Cell 184, 5506–5526, doi:10.1016/j.cell.2021.09.034 (2021).

2 Ablasser, A. et al. cGAS produces a 2’-5’-linked cyclic dinucleotide second messenger that activates STING. Nature 498, 380–384, doi:10.1038/nature12306 (2013).

3 Ishikawa, H., Ma, Z. & Barber, G. N. STING regulates intracellular DNA-mediated, type I interferon-dependent innate immunity. Nature 461, 788–792, doi:10.1038/nature08476 (2009).

4 Liu, S. et al. Phosphorylation of innate immune adaptor proteins MAVS, STING, and TRIF induces IRF3 activation. Science 347, aaa2630, doi:10.1126/science.aaa2630 (2015).

5 Sun, L., Wu, J., Du, F., Chen, X. & Chen, Z. J. Cyclic GMP-AMP synthase is a cytosolic DNA sensor that activates the type I interferon pathway. Science 339, 786–791, doi:10.1126/science.1232458 (2013).

6 Gonugunta, V. K. et al. Trafficking-Mediated STING Degradation Requires Sorting to Acidified Endolysosomes and Can Be Targeted to Enhance Anti-tumor Response. Cell Rep 21, 3234–3242, doi:10.1016/j.celrep.2017.11.061 (2017).

7 Burleigh, K. et al. Human DNA-PK activates a STING-independent DNA sensing pathway. Sci Immunol 5, doi:10.1126/sciimmunol.aba4219 (2020).

8 Taffoni, C. et al. DNA damage repair kinase DNA-PK and cGAS synergize to induce cancer-related inflammation in glioblastoma. EMBO J 42, e111961, doi:10.15252/embj.2022111961 (2023).

9 Guerra, J. et al. Lysyl-tRNA synthetase produces diadenosine tetraphosphate to curb STING-dependent inflammation. Sci Adv 6, eaax3333, doi:10.1126/sciadv.aax3333 (2020).

10 Fang, L., Ying, S., Xu, X. & Wu, D. TREX1 cytosolic DNA degradation correlates with autoimmune disease and cancer immunity. Clin Exp Immunol 211, 193–207, doi:10.1093/cei/uxad017 (2023).

11 Decout, A., Katz, J. D., Venkatraman, S. & Ablasser, A. The cGAS-STING pathway as a therapeutic target in inflammatory diseases. Nat Rev Immunol 21, 548–569, doi:10.1038/s41577-021-00524-z (2021).

12 Sharma, M., Rajendrarao, S., Shahani, N., Ramirez-Jarquin, U. N. & Subramaniam, S. Cyclic GMP-AMP synthase promotes the inflammatory and autophagy responses in Huntington disease. Proc Natl Acad Sci U S A 117, 15989–15999, doi:10.1073/pnas.2002144117 (2020).

13 Sliter, D. A. et al. Parkin and PINK1 mitigate STING-induced inflammation. Nature 561, 258–262, doi:10.1038/s41586-018-0448-9 (2018).

14 Yu, C. H. et al. TDP-43 Triggers Mitochondrial DNA Release via mPTP to Activate cGAS/STING in ALS. Cell 183, 636–649 e618, doi:10.1016/j.cell.2020.09.020 (2020).

15 Chahrour, M. & Zoghbi, H. Y. The story of Rett syndrome: from clinic to neurobiology. Neuron 56, 422–437, doi:10.1016/j.neuron.2007.10.001 (2007).

16 Guy, J., Gan, J., Selfridge, J., Cobb, S. & Bird, A. Reversal of neurological defects in a mouse model of Rett syndrome. Science 315, 1143–1147, doi:10.1126/science.1138389 (2007).

17 Cortelazzo, A. et al. Subclinical inflammatory status in Rett syndrome. Mediators Inflamm 2014, 480980, doi:10.1155/2014/480980 (2014).

18 van der Vaart, M. et al. Mecp2 regulates tnfa during zebrafish embryonic development and acute inflammation. Dis Model Mech 10, 1439–1451, doi:10.1242/dmm.026922 (2017).

19 Zalosnik, M. I., Fabio, M. C., Bertoldi, M. L., Castanares, C. N. & Degano, A. L. MeCP2 deficiency exacerbates the neuroinflammatory setting and autoreactive response during an autoimmune challenge. Sci Rep 11, 10997, doi:10.1038/s41598-021-90517-8 (2021).

20 Cronk, J. C. et al. Methyl-CpG Binding Protein 2 Regulates Microglia and Macrophage Gene Expression in Response to Inflammatory Stimuli. Immunity 42, 679–691, doi:10.1016/j.immuni.2015.03.013 (2015).

21 De Felice, C. et al. Rett syndrome: An autoimmune disease? Autoimmun Rev 15, 411–416, doi:10.1016/j.autrev.2016.01.011 (2016).

22 Leoncini, S. et al. Cytokine Dysregulation in MECP2-and CDKL5-Related Rett Syndrome: Relationships with Aberrant Redox Homeostasis, Inflammation, and omega-3 PUFAs. Oxid Med Cell Longev 2015, 421624, doi:10.1155/2015/421624 (2015).

23 Nan, X., Campoy, F. J. & Bird, A. MeCP2 is a transcriptional repressor with abundant binding sites in genomic chromatin. Cell 88, 471–481, doi:10.1016/s0092-8674(00)81887-5 (1997).

24 Jones, P. L. et al. Methylated DNA and MeCP2 recruit histone deacetylase to repress transcription. Nat Genet 19, 187–191, doi:10.1038/561 (1998).

25 Chen, L. et al. MeCP2 binds to non-CG methylated DNA as neurons mature, influencing transcription and the timing of onset for Rett syndrome. Proc Natl Acad Sci U S A 112, 5509–5514, doi:10.1073/pnas.1505909112 (2015).

26 Hansen, J. C., Ghosh, R. P. & Woodcock, C. L. Binding of the Rett syndrome protein, MeCP2, to methylated and unmethylated DNA and chromatin. IUBMB Life 62, 732–738, doi:10.1002/iub.386 (2010).

27 Tillotson, R. et al. Neuronal non-CG methylation is an essential target for MeCP2 function. Mol Cell 81, 1260–1275 e1212, doi:10.1016/j.molcel.2021.01.011 (2021).

28 Muotri, A. R. et al. L1 retrotransposition in neurons is modulated by MeCP2. Nature 468, 443–446, doi:10.1038/nature09544 (2010).

29 Bregnard, C. et al. Upregulated LINE-1 Activity in the Fanconi Anemia Cancer Susceptibility Syndrome Leads to Spontaneous Pro-inflammatory Cytokine Production. EBioMedicine 8, 184–194, doi:10.1016/j.ebiom.2016.05.005 (2016).

30 Gulmez Karaca, K., Brito, D. V. C., Zeuch, B. & Oliveira, A. M. M. Adult hippocampal MeCP2 preserves the genomic responsiveness to learning required for long-term memory formation. Neurobiol Learn Mem 149, 84–97, doi:10.1016/j.nlm.2018.02.010 (2018).

31 Nakatani, Y. & Ogryzko, V. Immunoaffinity purification of mammalian protein complexes. Methods Enzymol 370, 430–444, doi:10.1016/S0076-6879(03)70037-8 (2003).

32 Chamma, H., Guha, S., Laguette, N. & Vila, I. K. Protocol to induce and assess cGAS-STING pathway activation in vitro. STAR Protoc 3, 101384, doi:10.1016/j.xpro.2022.101384 (2022).

33 Schindelin, J. et al. Fiji: an open-source platform for biological-image analysis. Nat Methods 9, 676–682, doi:10.1038/nmeth.2019 (2012).

34 Krueger, F. Trim galore: a wrapper tool around Cutadapt and FastQC to consistently apply quality and adapter trimming to FastQ files., doi:10.5281/zenodo.7598955 (2015).

35 Kim, D., Paggi, J. M., Park, C., Bennett, C. & Salzberg, S. L. Graph-based genome alignment and genotyping with HISAT2 and HISAT-genotype. Nat Biotechnol 37, 907–915, doi:10.1038/s41587-019-0201-4 (2019).

36 Danecek, P. et al. Twelve years of SAMtools and BCFtools. Gigascience 10, doi:10.1093/gigascience/giab008 (2021).

37 Putri, G. H., Anders, S., Pyl, P. T., Pimanda, J. E. & Zanini, F. Analysing high-throughput sequencing data in Python with HTSeq 2.0. Bioinformatics 38, 2943–2945, doi:10.1093/bioinformatics/btac166 (2022).

38 Love, M. I., Huber, W. & Anders, S. Moderated estimation of fold change and dispersion for RNA-seq data with DESeq2. Genome Biol 15, 550, doi:10.1186/s13059-014-0550-8 (2014).

39 Ramirez, F. et al. deepTools2: a next generation web server for deep-sequencing data analysis. Nucleic Acids Res 44, W160–165, doi:10.1093/nar/gkw257 (2016).

40 Bendall, M. L. et al. Telescope: Characterization of the retrotranscriptome by accurate estimation of transposable element expression. PLoS Comput Biol 15, e1006453, doi:10.1371/journal.pcbi.1006453 (2019).

41 Mosley, S. R. & Baker, K. Isolation of endogenous cytosolic DNA from cultured cells. STAR Protoc 3, 101165, doi:10.1016/j.xpro.2022.101165 (2022).

42 Li, H. Minimap2: pairwise alignment for nucleotide sequences. Bioinformatics 34, 3094–3100, doi:10.1093/bioinformatics/bty191 (2018).

43 Sun, H. et al. A Nuclear Export Signal Is Required for cGAS to Sense Cytosolic DNA. Cell Rep 34, 108586, doi:10.1016/j.celrep.2020.108586 (2021).

44 Kranzusch, P. J., Lee, A. S., Berger, J. M. & Doudna, J. A. Structure of human cGAS reveals a conserved family of second-messenger enzymes in innate immunity. Cell Rep 3, 1362–1368, doi:10.1016/j.celrep.2013.05.008 (2013).

45 Stetson, D. B., Ko, J. S., Heidmann, T. & Medzhitov, R. Trex1 prevents cell-intrinsic initiation of autoimmunity. Cell 134, 587–598, doi:10.1016/j.cell.2008.06.032 (2008).

46 Beck-Engeser, G. B., Eilat, D. & Wabl, M. An autoimmune disease prevented by anti-retroviral drugs. Retrovirology 8, 91, doi:10.1186/1742-4690-8-91 (2011).

47 Marano, D., Fioriniello, S., D’Esposito, M. & Della Ragione, F. Transcriptomic and Epigenomic Landscape in Rett Syndrome. Biomolecules 11, doi:10.3390/biom11070967 (2021).

48 Yu, F., Zingler, N., Schumann, G. & Stratling, W. H. Methyl-CpG-binding protein 2 represses LINE-1 expression and retrotransposition but not Alu transcription. Nucleic Acids Res 29, 4493–4501, doi:10.1093/nar/29.21.4493 (2001).

49 Zhao, B. et al. Somatic LINE-1 retrotransposition in cortical neurons and non-brain tissues of Rett patients and healthy individuals. PLoS Genet 15, e1008043, doi:10.1371/journal.pgen.1008043 (2019).

50 Simon, M. et al. LINE1 Derepression in Aged Wild-Type and SIRT6-Deficient Mice Drives Inflammation. Cell Metab 29, 871–885 e875, doi:10.1016/j.cmet.2019.02.014 (2019).

51 Rice, G. I. et al. Reverse-Transcriptase Inhibitors in the Aicardi-Goutieres Syndrome. N Engl J Med 379, 2275–2277, doi:10.1056/NEJMc1810983 (2018).

52 Rookhuizen, D. C. et al. Induction of transposable element expression is central to innate sensing. bioRxiv, 2021.2009.2010.457789, doi:10.1101/2021.09.10.457789 (2021).

53 Ni, G., Ma, Z. & Damania, B. cGAS and STING: At the intersection of DNA and RNA virus-sensing networks. PLoS Pathog 14, e1007148, doi:10.1371/journal.ppat.1007148 (2018).

54 Kyle, S. M., Saha, P. K., Brown, H. M., Chan, L. C. & Justice, M. J. MeCP2 co-ordinates liver lipid metabolism with the NCoR1/HDAC3 corepressor complex. Hum Mol Genet 25, 3029–3041, doi:10.1093/hmg/ddw156 (2016).

55 Vila, I. K. et al. STING orchestrates the crosstalk between polyunsaturated fatty acid metabolism and inflammatory responses. Cell Metab 34, 125–139 e128, doi:10.1016/j.cmet.2021.12.007 (2022).

