## Supplementary figures for "The methyl-CpG-binding protein 2 inhibits cGAS-associated signaling"

Fig. S1 – S6

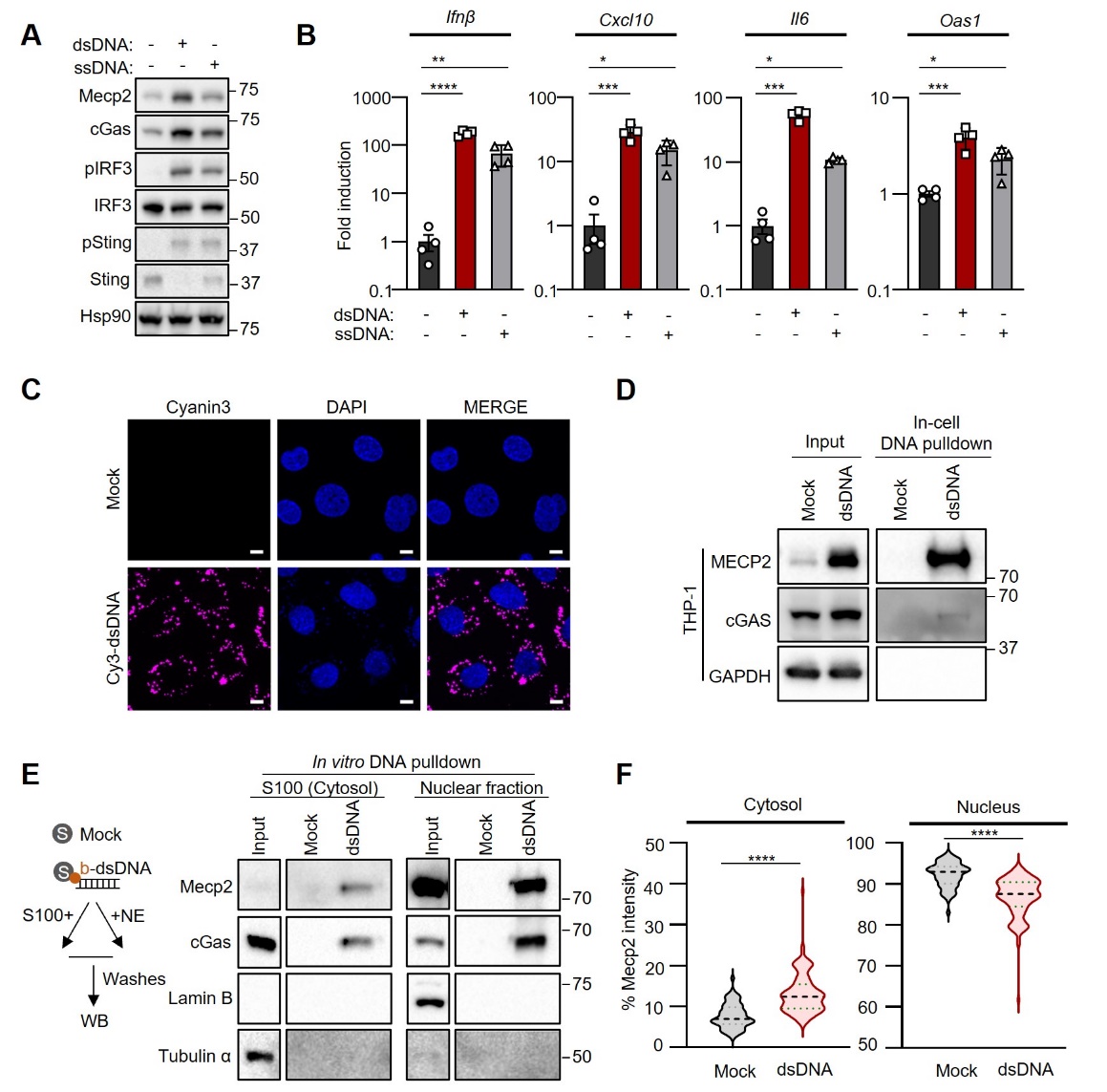

**Figure S1. Mecp2 interacts with cytosolic dsDNA**. **A** Whole cell extracts (WCE) prepared from WT-MEF transfected or not with dsDNA or ssDNA for 6 hours were analyzed by western blot (WB) using indicated antibodies. **B** *Ifnβ*, *Cxcl10*, *Il6*, *Oas1,* and *Mecp2* mRNA levels were analyzed in WT-MEF treated as in **A**. Graphs present the mean ± standard deviation of mean (SEM) from 4 independent experiments. **C** Imaging of WT-MEF cells transfected or not for 6 hours with Cy3-dsDNA and DAPI nuclear staining. Scale bar: 10µm. Images are representative of 3 independent experiments. **D** THP-1 were transfected or not with biotinylated b-dsDNA before whole-cell extract preparation and pull-down using streptavidin-affinity beads. Input and eluates were analyzed by WB using the indicated antibodies. **E** Left: Experimental scheme. Right: Cytosolic (S100) and nuclear (NE) fractions prepared from WT-MEF, were incubated with streptavidin beads alone or with streptavidin bead-bound b-dsDNA prior to pulldown. Input and eluates were analyzed by WB using the indicated antibodies. **F** Violin plots show the % of Mecp2 intensity in the cytosol in cells in experiments described in figure 1E (n=123 Mock transfected cells and 120 dsDNA-transfected cells).

WB are representative of 3 independent experiments. Significance was assessed using Student T-test. ns: non-significant. *P < 0.05, **P < 0.01, ***P < 0.001

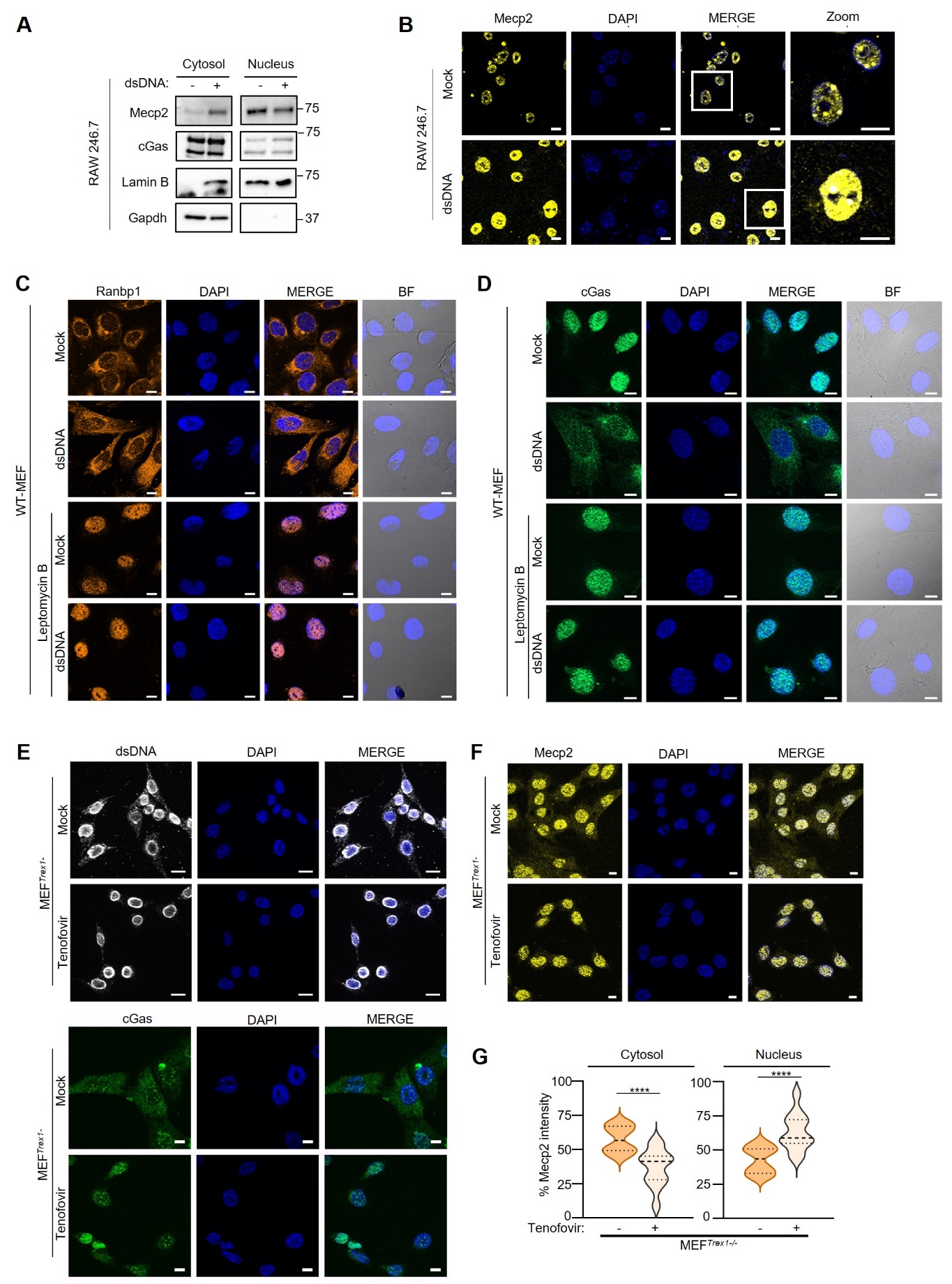

**Figure S2. dsDNA challenge triggers Mecp2 export.** **A** Cytosolic and nuclear extracts were prepared from WT-RAW264.7 transfected or not with dsDNA for 6 hours. Fractions were analyzed by WB using indicated antibodies. WB are representative of at least 3 independent experiments. **B** Immunofluorescence analysis was performed on WT-RAW264.7 treated as in **A** using anti-Mecp2 antibody and DAPI nuclear staining. Scale bar: 10µm. Images are representative of at least 3 independent experiments. **C** Immunofluorescence analysis was performed on WT-MEF treated or not with 20 nM of Leptomycin B for 1 hour prior to dsDNA transfection for 3 hours, using an anti-Ranbp1 antibody and DAPI nuclear staining. Scale bar: 10µm. Images are representative of two independent experiments. **D** Immunofluorescence was performed on WT-MEFs treated as in **C** except that an anti-cGas antibody was used. **E** Immunofluorescence analysis was performed on MEF*^Trex1-/-^* treated or not with tenofovir, using anti-dsDNA or anti-cGas antibodies, and DAPI nuclear staining. Scale bar: 20 µm. Images are representative of two independent experiments. **F**, As in **E** except that an anti-Mecp2 antibody was used. Scale bar: 10 µm. Images are representative of two independent experiments. **G** Violin plots show the % of Mecp2 intensity in the cytosol and in the nucleus; n=12 cells per condition.

Significance was assessed using Student T-test. ns: non-significant. *P < 0.05, **P < 0.01, ***P < 0.001 and ****P < 0.0001.

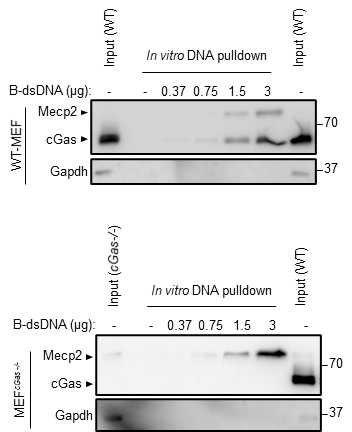

**Figure S3. Absence of cGas promotes enhanced Mecp2 interaction with dsDNA.** WCE prepared from WT-MEF (left) or MEF*^cGas-/-^* (right) were incubated with increasing quantity of streptavidin bead-bound b-dsDNA (as indicated). Input and eluates were analyzed by WB using the indicated antibodies.

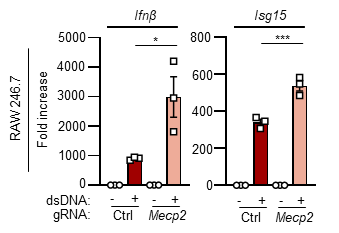

**Figure S4. Absence of Mecp2 enhances cGas-Sting activation.** RAW264.7^gCTRL^ or RAW264.7^gMecp2^ were challenged or not with dsDNA for 6 hours prior to gene expression analysis. Graph presents mean (± SEM) *Ifnβ*, and *Isg15* mRNA levels (n=3 independent experiments).

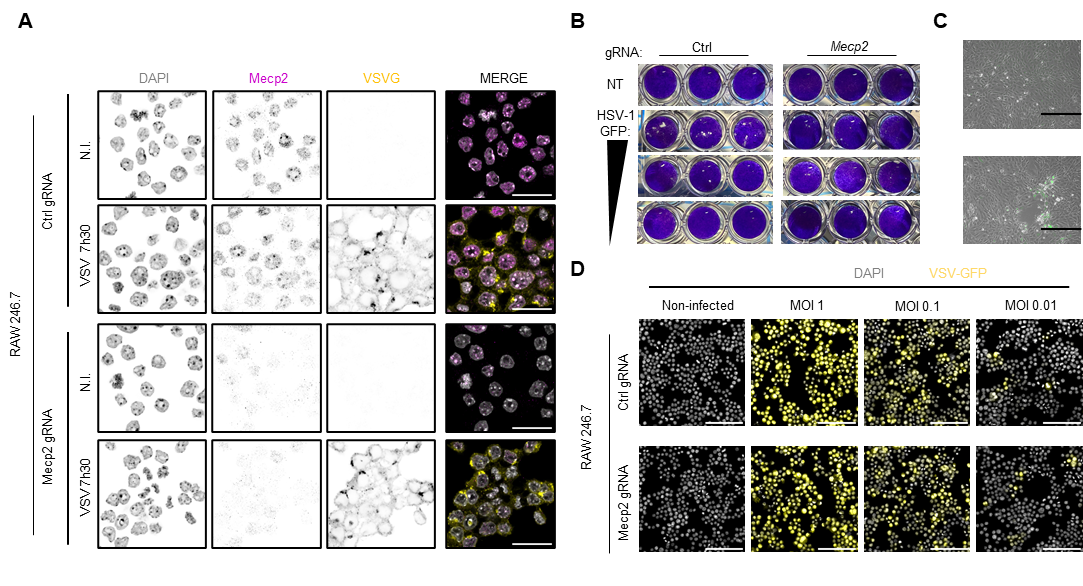

**Figure 5. Absence of Mecp2 enforces an antiviral state. A** RAW264.7^gCTRL^ or RAW264.7^gMecp2^ were infected or not with VSV for 7.5 hours prior to immunofluorescence analysis using anti-Mecp2 and VSV-G specific antibodies and DAPI nuclear staining. Images are representative of 2 independent experiments. Scale bar: 20µm. **B** Example images of plaques quantified in Figure 5E. **C** Representative images of MEF^gCTRL^ or MEF^gMecp2^ infected with HSV-1-GFP in Figure 5F. Scale bar: 400µm. **D** As in **A**, except that cells were infected with a range of VSV-G MOIs and GFP signal, attesting to VSV infection is shown. Images are representative of 2 independent experiments. Scale bar: 100µm.

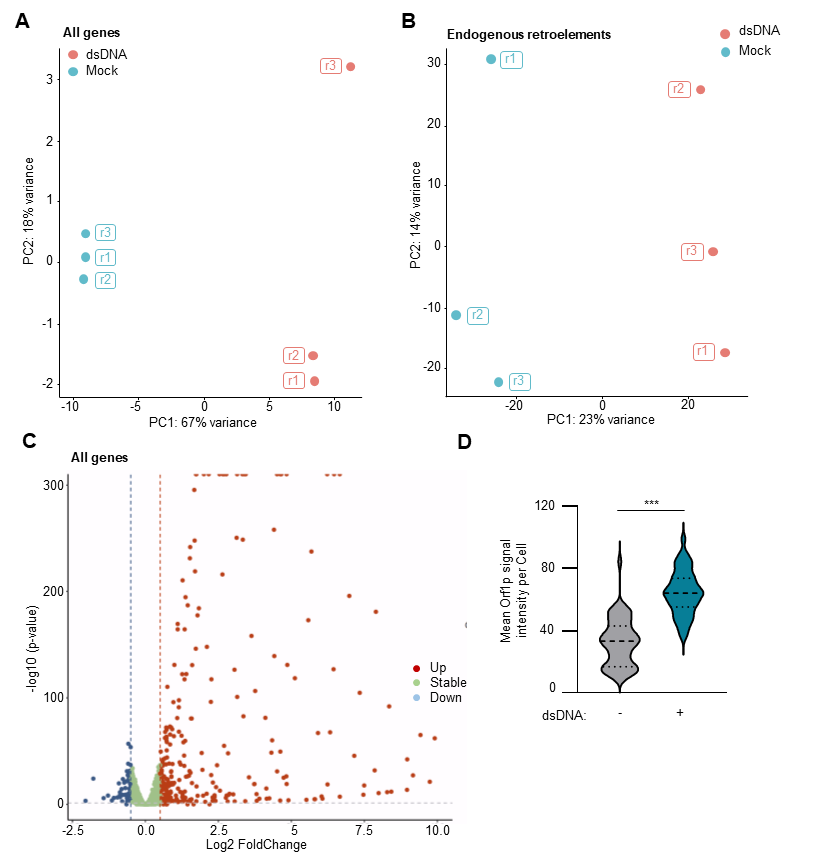

**Figure 6. Mecp2 deficiency leads to accumulation of immunogenic LINE-1 derived DNA. A** Principal Component Analysis (PCA) plot of all transcripts identified in RNAseq analyses conducted on RAW267.4 cells transfected of not with dsDNA for 6 hours. **B** PCA plot of transcripts corresponding to endogenous retroelements in samples from **A**. **C** Volcano plot representing upregulated, stable or downregulated transcripts in samples from **A**. **D** Violin plot shows the quantification of Orf1p signal in images acquired as in Figure 6D. n = 49 mock transfected cells; n= 87 dsDNA-transfected cells.

Significance was assessed using Student T-test. ***P < 0.001.
